# Genomic and phenotypic characterization of finger millet indicates a complex diversification history

**DOI:** 10.1101/2021.04.06.438618

**Authors:** Jon Bančič, Damaris A. Odeny, Henry F. Ojulong, Samuel M. Josiah, Jaap Buntjer, R. Chris Gaynor, Stephen P. Hoad, Gregor Gorjanc, Ian K. Dawson

## Abstract

Advances in sequencing technologies mean that insights into crop diversification aiding future breeding can now be explored in crops beyond major staples. For the first time, we use a genome assembly of finger millet, an allotetraploid orphan crop, to analyze DArTseq single nucleotide polymorphisms (SNPs) at the sub-genome level. A set of 8,778 SNPs and 13 agronomic traits characterizing a broad panel of 423 landrace accessions from Africa and Asia suggested the crop has undergone complex, context-specific diversification consistent with a long domestication history. Both Principal Component Analysis and Discriminant Analysis of Principal Components of SNPs indicated four groups of accessions that coincided with the principal geographic areas of finger millet cultivation. East Africa, the considered origin of the crop, appeared the least genetically diverse. A Principal Component Analysis of phenotypic data also indicated clear geographic differentiation, but different relationships among geographic areas than genomic data. Neighbour-joining trees of sub-genomes A and B showed different features which further supported the crop’s complex evolutionary history. Our genome-wide association study indicated only a small number of significant marker-trait associations. We applied then clustering to marker effects from a ridge regression model for each trait which revealed two clusters of different trait complexity, with days to flowering and threshing percentage among simple traits, and finger length and grain yield among more complex traits. Our study provides comprehensive new knowledge on the distribution of genomic and phenotypic variation in finger millet, supporting future breeding intra- and inter-regionally across its major cultivation range.

**Core ideas:**

- 8,778 SNPs and 13 agronomic traits characterized a panel of 423 finger millet landraces.
- 4 clusters of accessions coincided with major geographic areas of finger millet cultivation.
- A comparison of phenotypic and genomic data indicated a complex diversification history.
- This was confirmed by the analysis of allotetraploid finger millet’s separate sub-genomes.
- Comprehensive new knowledge for intra- and inter-regional breeding is provided.

## INTRODUCTION

Diversifying crop production is an important global objective to address human and environmental health concerns, such as malnutrition and the use of intensive unsustainable monoculture production systems (Bančič et al., 2021; von Grebmer et al., 2014). To achieve diversification, we need to increase focus on under-researched crops, also known as orphan crops. Some of these orphan crops are rich in micro- and macro-nutrients and can complement other crops in food production systems, even when farming conditions are adverse (Kamenya et al., 2021; Mustafa et al., 2019). Apart from their intrinsic value, many orphan crops have complex demographic histories that can shed light on broader crop domestication and diversification processes (Meyer & Purugganan, 2013). The high cost of generating genomic resources has however traditionally prevented genomic analyses for orphan crops, but a significant cost-reduction in high-throughput sequencing in the last decade has provided new opportunities for their research (Jamnadass et al., 2020).

Finger millet (*Eleusine coracana* (L.) Gaertn. subsp. *coracana*; Poaceae, subfamily Chloridoideae) is an annual small-grained cereal and an orphan crop with an essential role in smallholder food production systems in parts of Africa and Asia. The crop’s attractive characteristics include its rich nutritional profile, versatility in food usage, good storage properties, high market value, adaptability to poor production conditions, and flexibility in integrating into various farming approaches (Odeny et al., 2020; Sood et al., 2019). According to de Wet et al. (1984), the allotetraploid crop (*2n=4x*=36; genome constitution AABB, reported disomic inheritance) is thought to have been domesticated from wild *E. coracana* subsp. *africana* in either Uganda or the Ethiopian highlands of East Africa around five millennia ago. The domesticated subsp. *coracana* then spread to southern Africa while broadly maintaining sympatry with the subsp. *africana* wild form, and remaining in proximity to other *Eleusine* species (see a distribution of the eight *Eleusine* species in Africa depicted in **Fig. 1**). According to de Wet et al. (1984), the crop was introduced to India around three millennia ago, from where it was then dispersed further, including to Nepal. Therefore, East Africa is considered the primary centre of diversity and India an important secondary centre. The immediate wild progenitor of finger millet is believed to have arisen in East Africa by hybridization between two diploid species, the first of which being *E. indica*, which still occurs widely across Africa and is considered the (maternal) donor of what has become finger millet’s A sub-genome; and the second, still unknown (and now possibly extinct), diploid pre-B sub-genome donor (Liu et al., 2014; Zhang et al., 2019).

**Fig. 1:**
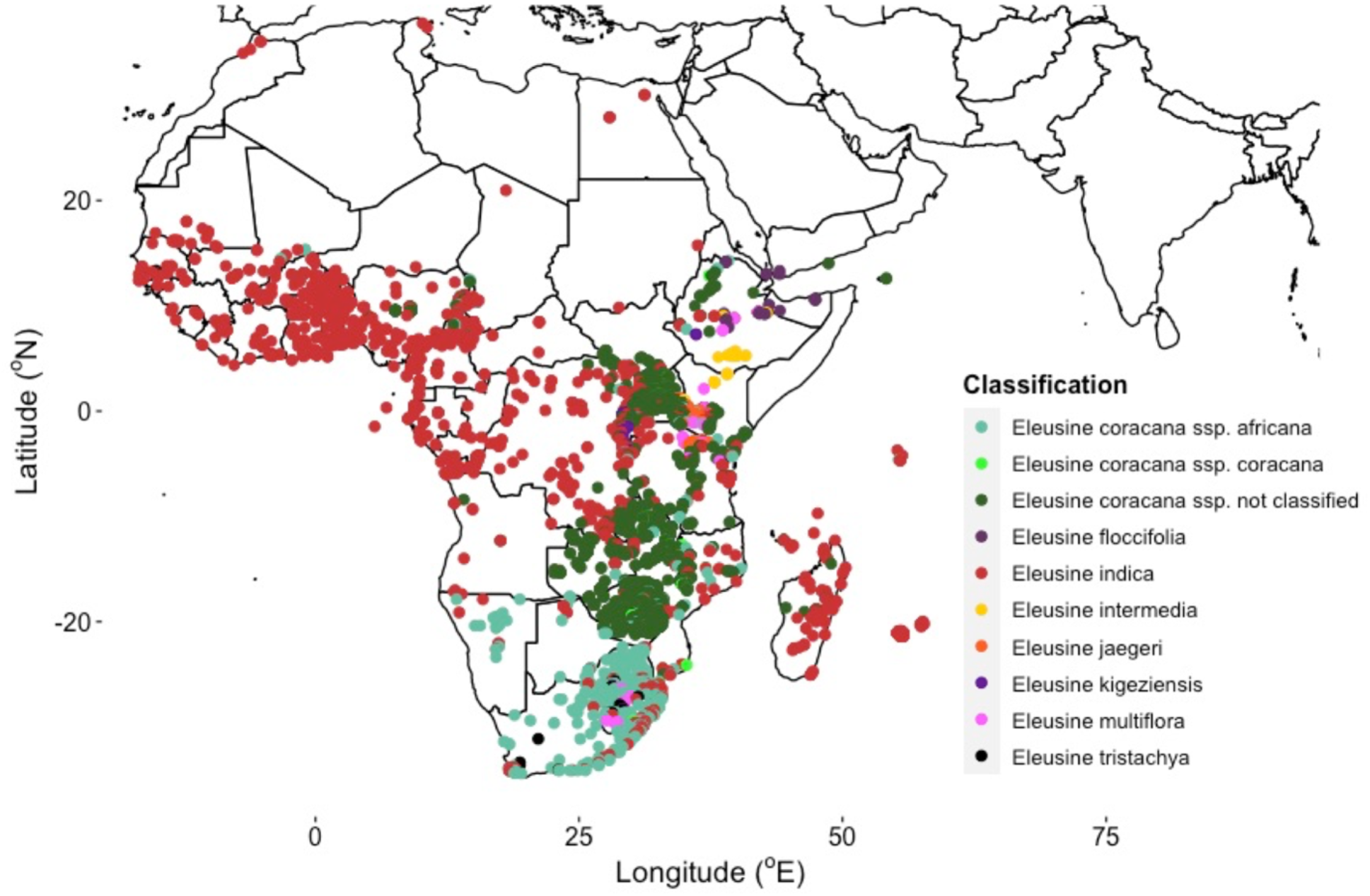
Distribution of eight *Eleusine* species across the African continent. A total of 9,250 known locations of *Eleusine* were extracted from the Global Biodiversity Information Facility (GBIF; https://www.gbif.org/). Some *Eleusine coracana* entries in GBIF were not classified at the subspecies level (‘not classified’ in key).

In Africa and Asia, subsistence farmers who rely on finger millet mostly grow landrace varieties, and systematic genetic improvement has been limited. This and its complicated biology reflect finger millet’s status as an orphan crop. The crop predominantly self-pollinates and has small flowers, which challenges artificial crosses (Dida & Devos, 2006). Moreover, its genomic status as an allotetraploid with an unknown B sub-genome donor has hampered the development of genomic resources. This is because difficulties in distinguishing between sub-genomes and thus the inaccurate calling of homologous versus homeologous single nucleotide polymorphism (SNP) positions are a possibility (Hatakeyama et al., 2018; Hittalmani et al., 2017). Landrace varieties however provide important opportunities to explore crop domestication and diversification. Many have been sampled for conservation in genebanks (Upadhyaya et al., 2006) and these accessions are available for use in breeding programs. They are also a resource to explore the crop’s diversification history, a topic that has so far received only limited attention.

The recent availability of a draft genome sequence and a robust linkage map for finger millet transform the potential for using genomic information to assist breeding and further understand the crop’s diversification history (Odeny et al., 2020). This paper uses a combination of genomic and phenotypic data to explore a broad panel of 423 finger millet landrace accessions sampled across its main cultivation regions in Africa and Asia. We make use of Diversity Arrays Technology sequencing (DArTseq) (Sansaloni et al., 2011) SNP markers that are, for the first time, placed onto finger millet sub-genomes using the recent genome assembly (https://phytozome-next.jgi.doe.gov/info/Ecoracana_v1_1). Our objective is to characterize the crop’s genomic and phenotypic variation to explore the diversification process and to provide insights for future breeding across its main cultivation range. We here explore multiple features of finger millet variation, including the geographic structuring of genomic and phenotypic diversity, sub-genome specific diversity profiles, germplasm migration events amongst geographic areas, and genetic architecture and selection patterns for agronomic traits. Our analysis provides essential information for the future development of finger millet and is a model for exploring other orphan crops.

## MATERIALS AND METHODS

### Plant material

This study uses the most extensive set of jointly genotyped and phenotyped finger millet landrace accessions to date from the crop’s main cultivation regions in Africa and South Asia. The panel initially contained 458 accessions (later reduced to 423 accessions for analysis) that as well as encompassing the main cultivation regions included a small number of accessions collected more widely (**Tab. S1**). In total, 19 accessions (designated as “other”) were collected outside the main cultivation regions or are of unknown origin. Our panel is from the ICRISAT genebank’s Core Collection of finger millet assembled by Upadhyaya et al. (2006). The 458 accessions are a subset with extensive phenotypic variation and have been the focus of breeder’s activities in recent years. Most of the Core Collection was initially sampled directly from farmers’ fields, although sometimes accessions were sampled from local markets (see, e.g., Rao, 1980).

### Collecting and processing genomic data

Leaf tissue was taken from single individuals of each of the 458 accessions, grown in a greenhouse at ICRISAT in Nairobi and dried with silica gel. Genomic DNA (gDNA) was extracted from finely-ground leaf material using the ISOLATE II Genomic DNA Kit (Bioline Pty Ltd) and according to the manufacturer’s instructions. The purity and quantity of extracted gDNA was determined by gel electrophoresis and a Qubit 2.0 Fluorometer (Life Technologies, Carlsbad, CA), respectively, with a final dilution of gDNA to 50 ng/μl. Genomic DNA was then delivered to the Integrated Genotyping Service and Support (IGSS) facility at the Bioscience eastern and central Africa-International Livestock Research Institute (BecA-ILRI) hub in Nairobi. This was for library construction and DArTseq data generation with SNP positions’ assignment using the v1.1 finger millet genome assembly (https://phytozome-next.jgi.doe.gov/info/Ecoracana_v1_1), and initial output quality control steps, using methods described previously (Sansaloni et al., 2011).

The IGSS facility generated 70,906 raw SNPs via DArTseq, which we then quality filtered using TASSEL (version 5.0; Bradbury et al. 2007) as illustrated in **Fig. S1**. The accessions and SNPs were filtered using a minimum call rate of 70%, which reduced our initial sample size of 458 accessions to 423 (**Tab. S1**). This constituted our final accession set for later data analyses. We then retained only those SNPs with a minor allele frequency (MAF) > 0.01. In further screening, we removed SNPs with a heterozygosity level greater than *2pq* × (1 – *F*), where *p* and *q* are the frequencies of the two allele states, and F is the inbreeding coefficient, for which a value of 0.5 was chosen due to the self-pollinating nature of finger millet. This last filter was applied to remove SNP calls that most likely originated from incorrectly collapsing homeologous positions across the two sub-genomes.

### Measuring linkage disequilibrium decay along chromosomes

Since our analysis is the first to use SNP marker positions assigned using a finger millet genome assembly, we initially explored associations among our high-quality SNP data set in our specific germplasm panel. We calculated linkage disequilibrium (LD) decay (r^2^) between all pairwise intra-chromosomal SNP combinations using a full correlation matrix either uncorrected or corrected for bias based on underlying genetic structure across the accessions. Calculations of *r*^2^ were performed in the R package *LDcorSV* (Mangin et al., 2012). Pairwise *r*^2^ values were then plotted against chromosomal physical distance. The decay curve was fitted based on Hill and Weir (1988) using R code from Marroni et al. (2011) and was then used to estimate the distance at which *r*^2^ decreased to 0.2. We estimated LD decay for each chromosome separately, for each sub-genome and for the genome as a whole.

### Genetic structure, differentiation and diversity

#### Genome-wide genetic structure

We used four approaches to characterize and visualize genetic structure. First, we used R packages *ade4* (Dray & Dufour, 2007) and *adegenet* (Jombart, 2008; the*find.clusters* function) to undertake Principal Component Analysis (PCA) and Discriminant Analysis of Principal Components (DAPC). For the latter, the optimum PC number and cluster number (*k*) were set based on the change in the curve shape of profiles (**Figs. S2**, **S3**). Second, we constructed a genomic relationship matrix (GRM) using a centred-identity-by-state method (Endelman & Jannink, 2012) in TASSEL. A heatmap of the GRM, calculated using the unweighted pair group method with arithmetic mean (UPGMA) cluster algorithm, was visualized using the *heatmap* R function (R Core Team, 2019). Third, we constructed using TASSEL and plotted with R package *phangorn* (Schliep, 2011) an unweighted neighbour-joining (NJ) tree (Saitou & Nei, 1987) of accessions. We extended NJ analysis to consider not only genome-wide markers but SNPs pooled at the sub-genome level. Fourth, we visualized the country-level geographic distribution of finger millet accessions assigned to their genetic clusters using R packages *rworldmap* (South, 2011) and *ggplot2* (Wickham, 2009).

#### Detection of introgression and gene flow

We used Treemix (Pickrell & Pritchard, 2012) to infer the most likely evolutionary history and evidence for introgression amongst groups of accessions assigned to four geographic areas of sampling that we term ‘East Africa’, ‘Southern Africa’, ‘India’ and ‘Nepal’ (**Tab. S1**). For simplification, the small number of accessions sampled from countries outside these locations (9 of all 404’ known’-location accessions) were assigned to the most proximate of our defined four areas (indicated with ellipses in **Fig. 2d**). The 19 accessions designated as “other” (see Plant material section) were excluded from geographic area analysis. **Table S1** provides complete information, but, in brief: the area ‘East Africa’ included a small number of additional accessions (N = 5) from Nigeria and Senegal; and ‘India’ included a few extra lines (N = 4) from Sri Lanka, the Maldives and Pakistan. Using Treemix, maximum-likelihood population trees were constructed based on genome-wide SNPs using blocks of 50 SNPs and ‘East Africa’ rooted as an out-group. The number of tested migration events was varied from zero to three. Bootstrap replicates were generated using 50 SNPs to evaluate the robustness of tree topology, and Treemix R plotting functions were used to visualize results.

**Fig. 2:**
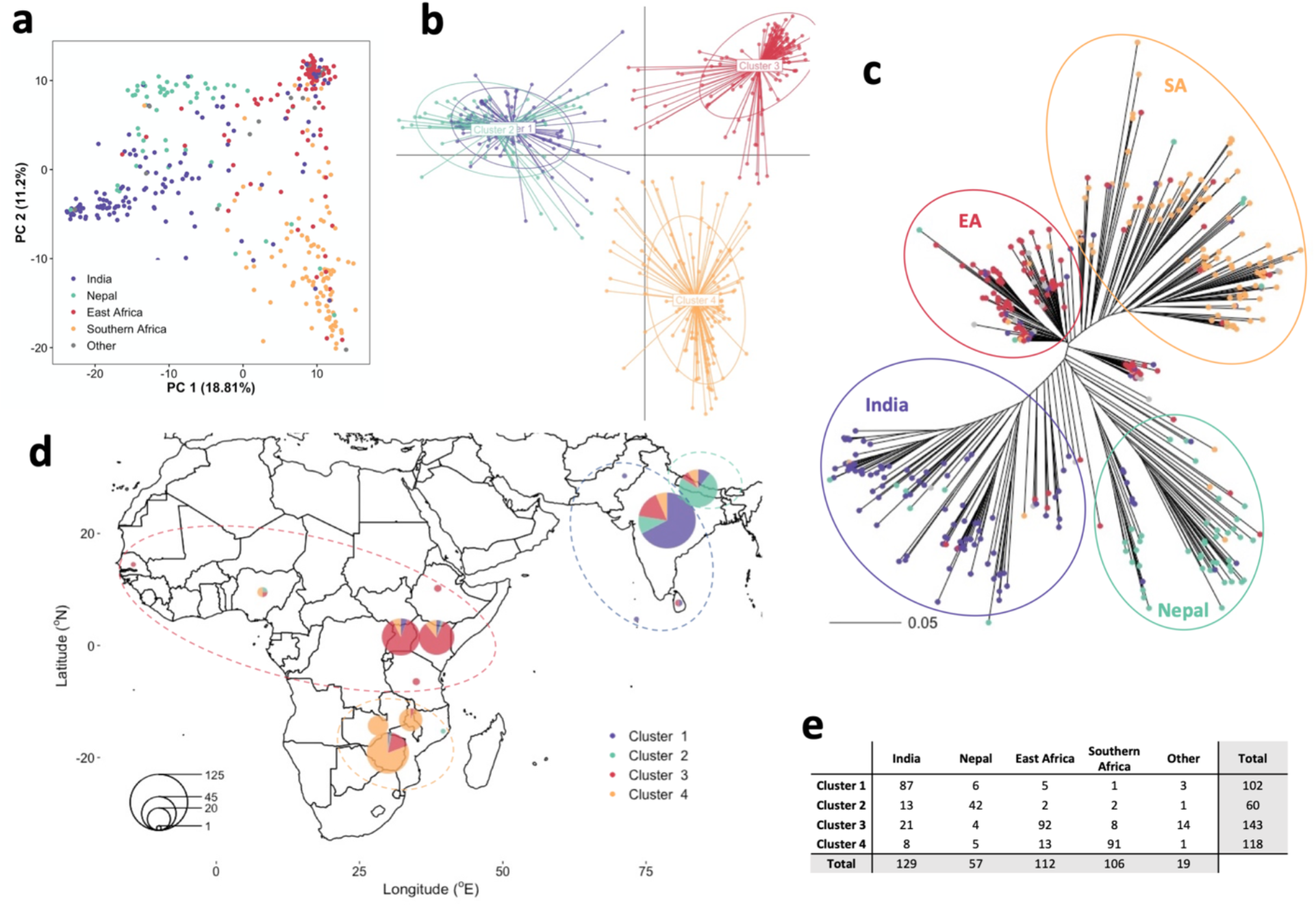
Geographical distribution and genetic structure of 423 finger millet accessions. **a**, Score plot using the first two principal components (PCs) illustrates the genetic structure of finger millet coloured according to geographic areas. **b**, Genetic clusters identified by DAPC and coloured according to their corresponding geographic area in **d**). **c**, A phylogenetic neighbourjoining tree with tips coloured by geographic area and enclosing circles representing generalized clades. **d**, Geographical distribution of accessions with pie chart size representing the number of accessions collected in a particular country and slice size representing the probability of belonging to a specific genetic cluster. Dotted ellipses indicate the extent of our applied geographic areas (drawing in small groups of outliers; see Materials and Methods and **Tab. S1** for further information on geographic area assignments). **e**, A table of accessions assigned to geographic areas and genetic clusters; accessions without known location are designated as ‘Other’.

#### Chromosome-level nucleotide diversity and differentiation

We analyzed diversity and differentiation at a chromosome level for both sub-genomes of finger millet for accessions assigned to four geographic areas of sampling (areas as explained above). For each area, we calculated ***π*** (Nei and Li, 1979) as our estimator of diversity and pairwise F_ST_ values (Weir and Cockerham, 1984) as our estimator of differentiation, using VCFtools (Danecek et al., 2011) and a 1 Mb non-overlapping sliding window. Genetic differentiation estimates were calculated for all six possible pairwise combinations of geographic areas. Results were plotted against chromosomal positions using R package *ggplot2*.

#### Summarized gene diversity and differentiation statistics

We summarised genome-wide gene diversity and differentiation for the four geographic areas. Genome-wide gene diversity (H; Nei, 1973) and pairwise F_ST_ values based on all high-quality SNPs were calculated with R package *hierfstat* (Goudet, 2005). We also computed H values for the four defined genetic clusters (see above) of finger millet accessions (these approximate our four defined geographic areas, as will be explained below). We further extended our analysis of geographic areas to consider genome-wide markers and diversity at the sub-genome level. We also used R package *pegas* (Paradis, 2010) for an analysis of molecular variance (AMOVA) that partitioned genetic variation within and among our geographic areas (or clusters) as part of the total panel, based on 100,000 permutations.

### Collecting and analyzing phenotypic data

To collect information on phenotypic variation in finger millet, we characterized our initial panel of 458 accessions for 13 life history and other traits (**Tab. S2**) in a field trial that used a complete randomized block design with two replications. The trial was conducted over the long rainy season of 2015 at the Kenya Agricultural and Livestock Research Organization field station at Kiboko in Eastern Kenya (coordinates: 2° 20’ N, 37° 45’ E; altitude: 960m; annual temperatures: min. 16.6°C, max. 29.4°C, average 23.0°; sandy clay loam calcareous soil). Seeds of each accession were drilled in 2 m long plots of two rows spaced 50 cm apart. Within rows, plants were thinned after establishment to a spacing of 10 cm. Five plants for each plot were randomly selected for data recording, though some traits were recorded on a whole plot basis (**Tab. S2**). The traits we measured included features of morphology, physiology and yield that are important for crop production and the integration of finger millet in mixed crop farming systems (Dawson et al., 2019), including in intercrop systems (Brooker et al., 2015), that are of particular interest in breeding research (Bančič et al., 2021).

Our analysis of variance of phenotypic data used the following statistical model:

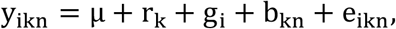

where *y_ikn_* is generally the average phenotype of five plants of the *i*^th^ accession tested in the *k*^th^ replication (*k* = 1, 2) and in the *n*^th^ block (*n* = 1,…, n); *μ* is the intercept; *r_k_* is the fixed effect of a replication; *g_i_* is the random accession (genotype) effect assuming 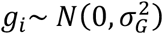; *b_kn_* is the random block effect assuming 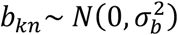; and *e_ikn_* is the random residual assuming 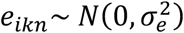. Here *N*(.,.) denotes a normal random variable and 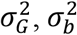 and 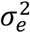 are respectively variances between accessions, blocks and residuals. We treated accessions as fixed effects to obtain best linear unbiased estimates (BLUEs) for each phenotypic trait. Models were fitted using R package *ASReml-R* (version 4.1.0.90; Butler et al. 2017). We estimated broad-sense heritability across the two replications of the trial (*p* = 2) for each trait as 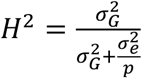. Distributions of BLUEs for each phenotypic trait and Pearson’s correlation coefficients (*r*; Pearson, 1895) between all pairwise combinations were calculated and visualized using R (R Core Team, 2019). The coefficient of variation (CV) for each phenotypic trait was calculated as 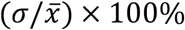 using BLUEs.

We then performed Principal Component Analysis using BLUEs of traits to understand the phenotypic structure in these data using R package *factoextra* (Kassambara & Mundt, 2020). Variation of BLUEs within our defined geographic areas and genetic clusters was also visualized using boxplots.

### Trait genetic architecture and further genomic-phenotypic comparison

#### Genome-wide association analysis

To test each SNP’s effect on the 13 phenotypic traits, we ran a genome-wide association study (GWAS) in TASSEL using a linear mixed model that corrected for genetic structure and cryptic relatedness (Yu et al., 2006). BLUEs calculated for each trait were taken as phenotype, and 8,778 SNPs were taken as genotype. The model estimated variance components only once using the P3D method. A Bonferroni threshold of 5% was applied to account for multiple testing to declare significant marker-trait associations (MTAs). Manhattan and quantile-quantile (Q-Q) plots of the GWAS results were visualized with R package *qqman* (Turner, 2014). Putative candidate genes were selected using a 25 kb genomic interval both upstream and downstream of a significant MTA. The interval was queried against the current finger millet genome assembly using the Integrated Genomics Viewer (IGV, v. 2.82; Robinson et al., 2011).

#### Clustering SNP effects

To further evaluate the genetic architecture of the 13 phenotypic traits, we explored the distributions of SNP effects from a ridge regression model (RR-BLUP) (Kooke et al., 2016). BLUEs calculated for each trait scaled by their standard deviation were taken as phenotype, and an imputed 8,778 SNP matrix were taken as genotype. A *k*-nearest neighbour imputation of the SNP matrix was performed in TASSEL using default parameters and RR-BLUP models were fitted in R package *AlphaMME* (https://github.com/gaynorr/AlphaMME). SNP effects from the RR-BLUP model were then used to: first, calculate pairwise Euclidean distances over the first five mathematical moments in R package *moments* (v0.14; Komsta and Novomestky, 2011); and, second, construct a dendrogram from the Euclidean distance matrix using the UPGMA cluster algorithm with the *hclust* R function (R Core Team, 2019).

#### Comparison of phenotypic and genome-wide gene diversity by geographic area

A simple yet useful way to shed light on the particular selection histories of crops as they take different diversification pathways within specific geographic contexts is to compare phenotypic diversity levels with levels of underlying genome-wide gene diversity. Here, we adopt a straightforward approach for initial comparisons that involves individual phenotypic trait CV values and gene diversity (H) values for genome-wide SNP data. We calculate CV/H as the comparator for each of our four defined geographic areas to check primarily for rank differences across areas that may indicate particular phenotypic selection pressures by area. This type of approach is exemplified classically in studies of the diversification of tomato, where high phenotypic variation observed at specific traits is accompanied by overall underlying genomic diversity bottlenecks (Rodríguez et al., 2011).

## RESULTS

### Genome-wide SNP data cover the entire finger millet chromosome complement but occur at a higher density in the A sub-genome

The IGSS facility generated 70,906 raw SNPs, which were filtered into a final marker set of 8,778 SNPs for 423 accessions for subsequent analyses. Eight thousand and ninety-six (8,096) SNPs out of the 8,778 had been previously mapped to chromosomes (**Tab. S3**). Initial analysis showed that the full complement of finger millet chromosomes was covered but that chromosome-level SNP density was higher for the A sub-genome (**Tab. 1**; approx. twice the density of markers of the B sub-genome). Across both sub-genomes, the mean SNP density was ~8.6 per Mb. Consistent with expectations of significantly lower recombination in central chromosomic regions of selfing cereals (e.g., Bustos-Korts et al., 2019); SNP density was generally considerably higher toward the ends of chromosomes (**Figs. S4**, **S9c**). The proportion of markers removed from our initial raw data due to likely being homeologous was a relatively high 3.9% (2,733 out of 70,906; markers above the red curve in **Fig. S1e**). Therefore, precautions with homeologous markers are indicated for finger millet. The observed heterozygosity for the markers in our final set of 8,778 SNPs was very low (median 0.048 and mean 0.057; **Tab. S3**), indicating that most finger millet accessions are highly inbred.

**Tab. 1.** Distribution of SNPs by finger millet chromosomes. SNP density per chromosome was calculated for individual 1 Mb non-overlapping windows as shown in **Figs. S4** and **S9c**.

| Chromosome | Number of SNPs | Chromosome length (Mb) | SNP density (per Mb) |
| --- | --- | --- | --- |
| 1A | 742 | 57.13 | 12.4 |
| 2A | 683 | 60.90 | 11.1 |
| 3A | 570 | 53.87 | 10.9 |
| 4A | 618 | 39.68 | 15.4 |
| 5A | 1057 | 66.30 | 15.4 |
| 6A | 357 | 59.47 | 5.9 |
| 7A | 553 | 48.45 | 11.2 |
| 8A | 307 | 47.22 | 7.6 |
| 9A | 486 | 45.36 | 11.6 |
| **Sub-genome A** | **5,373** | **53.15** | **11.3** |
| 1B | 196 | 72.49 | 3.7 |
| 2B | 323 | 70.83 | 5.3 |
| 3B | 136 | 63.91 | 2.9 |
| 4B | 158 | 50.88 | 2.9 |
| 5B | 659 | 80.22 | 7.8 |
| 6B | 244 | 74.22 | 5.0 |
| 7B | 267 | 58.85 | 7.1 |
| 8B | 294 | 60.62 | 5.4 |
| 9B | 446 | 62.05 | 7.4 |
| **Sub-genome B** | **2,723** | **65.20** | **5.7** |
| **Scaffolds** | **682** |  |  |
| **Whole genome** | **8,778** | **1072.44** | **8.6** |

### Linkage disequilibrium decay along chromosomes shows expected patterns for both sub-genomes

Our work is the first that has used a finger millet genome assembly to assign physical SNP positions for a broad accession panel, so understanding patterns of LD in A and B sub-genomes is of particular relevance. Our calculations indicated the expected overall pattern of LD decay and the importance of correcting for underlying genetic structure among accessions. For the genome as a whole, *r*^2^ decayed to 0.2 at a distance of 106 kb when correcting for genetic structure (compared to a distance of 1.77 Mb for the naïve model). Decay was slower for the B sub-genome (*r^2^* = 0.2 at 168 kb, with correction) than the A sub-genome (*r^2^* = 0.2 at 88 kb), and varied markedly across chromosomes within sub-genomes (**Fig. S5**). Overall, the relatively slow rate of LD decay was consistent with other self-pollinating crops (Flint-Garcia et al., 2003), including other millets (Jaiswal et al., 2019).

### Genome-wide genetic structure reveals strong geographic differentiation

Our analysis of genome-wide genetic structure in finger millet revealed clear differentiation patterns (**Fig. 2**). Both PCA and DAPC analyses (DAPC using an optimum principal component number PC = 4 according to **Fig. S2** and an optimal cluster number *k* = 4 according to **Fig. S3**) revealed a genetic structure that corresponded with the four sampled geographic areas of finger millet cultivation (**Fig. 2a, b, d**). Accessions from Africa and Asia regions were separated along the first PC, which explained 18.8% of the total variation, while the second PC, which explained 11.2% of the total variation, discriminated between geographic areas within the two continents (**Fig. 2a**). The overlap between genetic clusters and geographic areas was particularly strong in Africa (**Fig. 2e**, **Tab. S4**). The clear overall geographic structuring of genetic variation, with a degree of admixture, was also evident in our NJ tree (**Fig. 2c**), on the map of sampled geographic areas showing accessions’ assignments to genetic cluster groups (**Fig. 2d**), and in a heatmap of GRM with UPGMA clustering (**Fig. S6**). The more extensive admixture of finger millet in Asia than Africa, most clearly observed in **Fig. 2d**, is consistent with more formal breeding of the crop in Asia, supported by cross-regional germplasm transfer. For example, in India, an improvement program initiated in the 1960s involved crosses between African and Asian accessions, resulting in ‘Indaf’ varieties (Mirza & Marla, 2019; see more in next section). In further NJ analysis that considered SNPs for sub-genomes A and B separately, clear geographic structuring of genetic variation was evident in both cases (**Fig. S7**).

### Genetic introgression and gene flow analysis suggests a migration event between East Africa and India

Our Treemix analysis of past admixture events in finger millet suggested a potential historic introgression from East Africa to India. Using ‘East Africa’ as an out-group, the maximumlikelihood tree without migration events (**Figs. 3a** and **S8a**), accounting for drift alone, corresponded to the phylogenetic tree (see **Fig. 2c**). When migration events were allowed, a single event from ‘East Africa’ to ‘India’ was inferred (**Figs. 3b** and **S8b**). These results are consistent with the cultivated finger millet’s demographic history as previously speculated by non-genomic analysis approaches (de Wet et al., 1984; Hilu & de Wet, 1976). These findings are also consistent with known breeding introductions from Africa to Asia (see previous section).

**Fig. 3:**
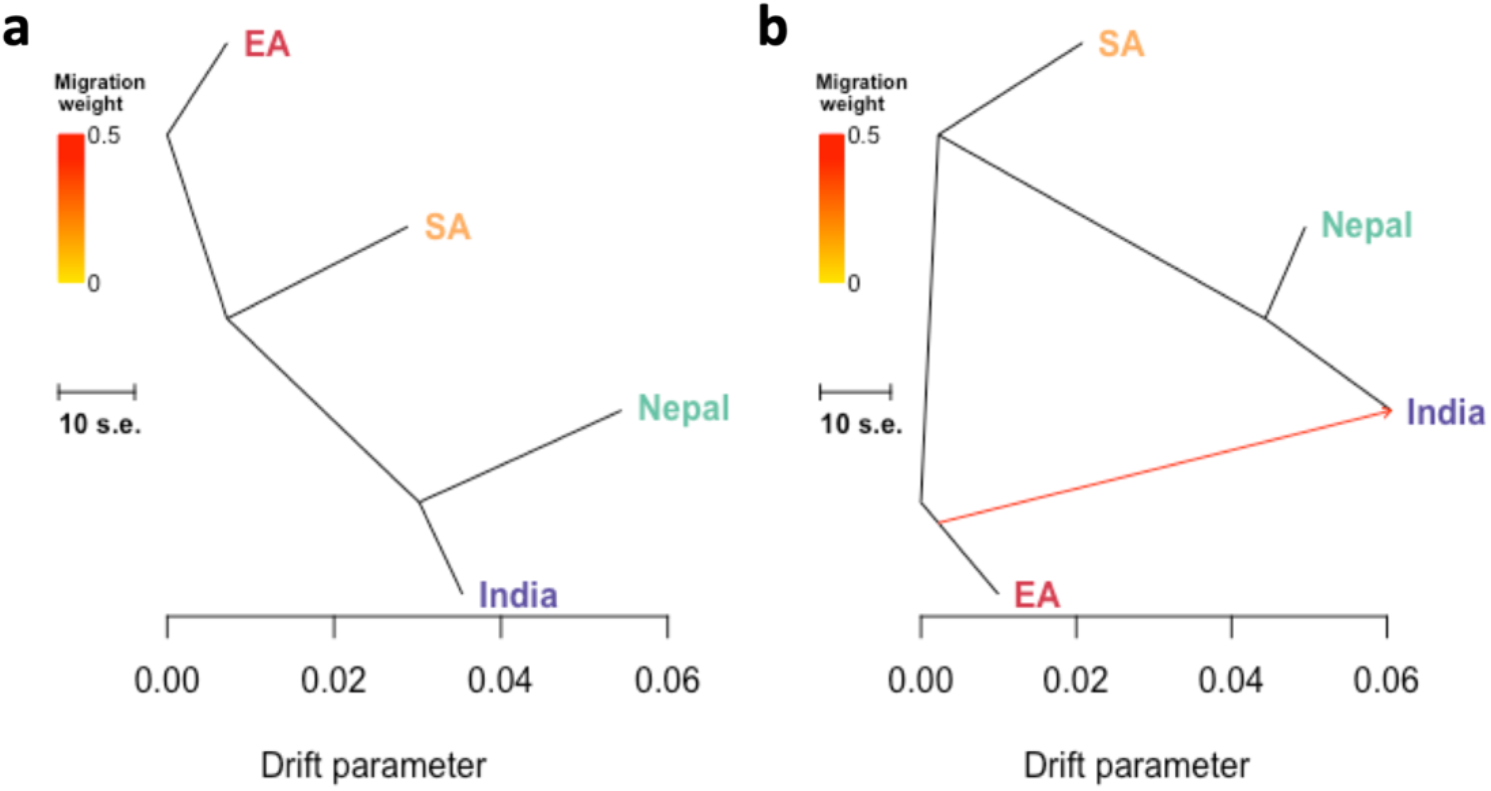
Inferred finger millet maximum-likelihood population trees with admixture events. The structure of the graphs inferred by Treemix for four geographic areas of finger millet. **a**, with no migration events. **b**, with one migration event. The migration arrow is coloured according to its weight and represents the fraction of ancestry derived from the migration edge. Horizontal branch lengths are proportional to the amount of genetic drift that has occurred on the branch. The scale bar shows ten times the average standard error (s.e.) of the accessions in the sample. The residual fit from the graph is shown in **Fig. S8**.

### Chromosome-level nucleotide diversity and differentiation profiles vary by region

Our analysis of nucleotide diversity (*π*) along chromosomes showed differences in relative diversity for geographic areas by chromosome, including for homeologous (A and B sub-genome) chromosomes (**Fig. S9a**). Notable was relatively high diversity along chromosome 1A for ‘Nepal’ (not seen on chromosome 1B) and along chromosomes 5B and 9B for ‘Southern Africa’ (not seen on chromosomes 5A and 9A). Pairwise chromosome-level F_ST_ values for geographic areas (**Fig. S9b**) reflected these different diversity profiles, again indicating chromosome-specific, subgenome-based differences. Of note was high differentiation between both of our African geographic areas and both our Asian geographic areas along chromosome 5A that was not replicated on chromosome 5B.

### Summarized gene diversity statistics confirm regional differentiation and sub-genomic differences

Our calculation of genome-wide gene diversity (H) values indicated that the ‘Nepal’ region contained the most diversity and the ‘East Africa’ region the least (**Tab. 2**). The ranking of diversity levels corresponded when the analysis was repeated based on genetic clusters that approximate geographic areas. Estimates were, as expected, overall lower when based on genetic clusters, as the most prominent within-area genetic admixture has in this case been reassigned (see **Fig. 2**).

**Tab. 2.** Characterization of gene diversity (H). Genome-wide measurement of finger millet gene diversity was undertaken for four geographic areas (left value) and corresponding genetic clusters identified by DAPC analysis (right value). The values presented for individual sub-genomes are for geographic areas only. The total sample size for genome-wide estimates varies because of the non-assignment of some accessions to geographic areas (see Materials and Methods). The area or cluster with highest diversity is underlined.

| Geographic area /<br>Genetic cluster | Sample<br>size | H |  |  |
| --- | --- | --- | --- | --- |
|  |  | Genome-wide | Sub-genome<br>A | Sub-genome<br>B |
| India / Cluster 1 | 129 / 102 | 0.259 / 0.215 | 0.293 | 0.212 |
| Nepal / Cluster 2 | 57 / 60 | <u>0.282</u> / <u>0.265</u> | <u>0.336</u> | 0.252 |
| EA / Cluster 3 | 112 / 143 | 0.199 / 0.179 | 0.200 | 0.208 |
| SA / Cluster 4 | 106 / 118 | 0.260 / 0.252 | 0.230 | <u>0.368</u> |

Pairwise F_ST_ values summarised for genome-wide SNPs (**Tab. S5**) confirmed our PCA and DAPC results (**Fig. 2a, b, d**), which revealed primary geography-based partitioning between Africa and Asia and then between areas within these regions. The highest differentiation was between ‘Southern Africa’ and ‘Nepal’ accessions (F_ST_ = 0.167). Our two AMOVA analyses revealed that 16% of total genome-wide variation partitioned among our four geographic areas and 24% among our corresponding but genetically-defined clusters (*p* ≤ 0.001 that no structuring in both cases).

In the case of gene diversity (H) calculations, we also analyzed geographic areas for separate A and B sub-genomes (**Tab. 2**). The ranking of diversity by geographic area varied for the two sub-genomes, with ‘Nepal’ (still, compared to the entire genome) ranking highest for the A subgenome but ‘Southern Africa’ highest for the B sub-genome; ‘East Africa’ consistently ranked lowest. The observed diversity ranking change is reflected in the relative spread of accessions from each geographic area in sub-genome NJ trees (**Fig. S7**). It appears to be based on changes in relative diversity for specific pairs of homeologous chromosomes (**Fig. S9a**), notably chromosomes 5B versus 5A, and 9B versus 9A (diversity relatively high at 5B and 9B for ‘Southern Africa’; see also above).

### Significant variation in phenotypic traits partitions by region

Our analysis of 13 phenotypic traits for 423 finger millet accessions revealed extensive variation (CV from 8.58 for threshing percentage to 48.25 for productive tiller number) and medium-to-high *H*^2^ values (from 0.35 for grain yield to 0.95 for days to flowering). The level of variation detected and heritability values were generally consistent with the crop’s previous field trials (e.g., Bharathi, 2011; Manyasa et al., 2016).

Summary statistics are presented in **Tab. S6**, and trait distributions and between-trait correlations in **Fig. S10**. The strongest positive correlation was between leaf length and plant height (*r* = 0.83, p < 0.001), and the strongest negative correlation between leaf width and the number of productive tillers (*r* = −0.64, p < 0.001). Considering the three key traits of grain yield, plant height and days to flowering, a medium-level positive correlation was observed between plant height and days to flowering (*r* = 0.50, p < 0.001), a moderate positive correlation between plant height and grain yield (*r* = 0.19, p < 0.001), and no correlation between days to flowering and grain yield (*r* < 0.01, NS). Principal component analysis using BLUEs of traits further illustrated the levels of correlation between different traits (**Fig. 4a**).

**Fig. 4:**
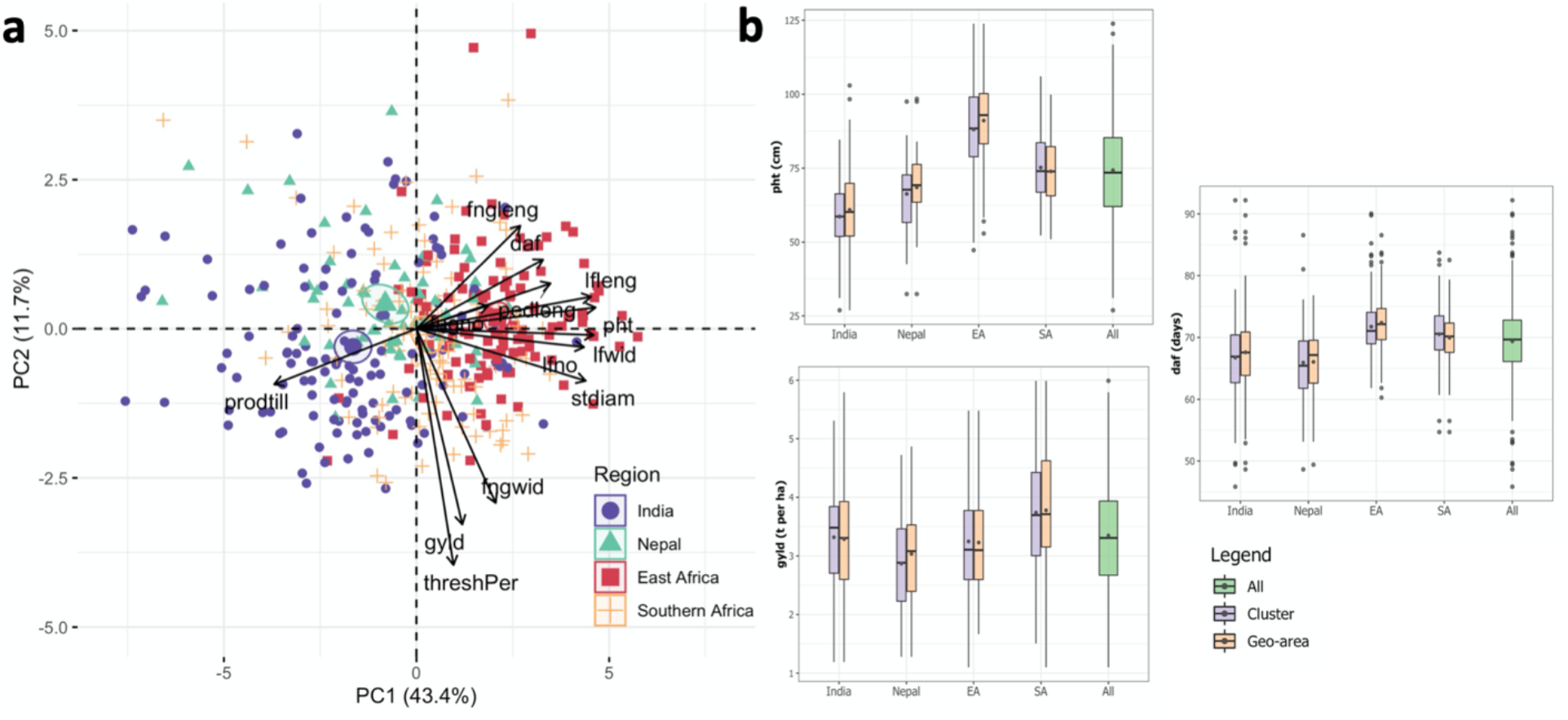
Summary of phenotypic data for 423 finger millet accessions. **a**, The Principal component analysis (PCA) biplot shows vectors of each of the 13 phenotypic traits/variables (black arrows) and the PCA scores for combined phenotypes of 404 accessions (represented by points) coloured according to geographic areas; scores of accessions (19) without known location were excluded. The magnitude of the vectors shows the strength of their contribution to each PC. Vectors pointing in similar directions indicate positively correlated traits, vectors pointing in opposite directions indicate negatively correlated variables, and vectors at approximately right angles indicate low or no correlation. Coloured ellipses represent 95% confidence intervals around the centroid (bold symbol) for each area. stdiam, stem diameter; lfleng, leaf length; lfwid, leaf width; lfno, leaf number; fngleng, finger length; fngwid, finger width; fngno, finger number; pedleng, peduncle length; pht, plant height; prodtill, production tillers; daf, days to flowering; thershPer, threshing percentage; gyld, grain yield. **b**, Boxplots of the statistical distribution of individual trait values divided according to genetic clusters identified by Discriminant Analysis of Principal Components (purple boxplots), geographic areas (orange boxplots) and the total phenotypic variation (green boxplots). The middle bar and the point inside each boxplot represent median and mean, respectively. Examples are given for days to plant height (pht) and grain yield (gyld). Results for the other 10 phenotypic traits are shown in **Fig. S11**.

Similar to genomic data, we used PCA scores of 404 accessions to examine our phenotypic data structure. Confidence ellipses based on geographic areas (**Fig. 4a**; confidence level set to 95% around area centroid) indicated a degree of separation along the first PC for all four geographic areas based on combined phenotypes, most clearly separating ‘India’ and ‘East Africa’. Accessions from ‘Nepal’ and ‘Southern Africa’ were relatively less differentiated compared to using genome-wide SNP data (see, e.g., F_ST_ values, **Tab. S5**). Trait-specific boxplots of phenotypic variation by region or genetic cluster (**Fig. 4b** and **Fig. S11**) illustrated a divergent pattern for some traits. For example, for plant height ‘India’ and ‘East Africa’ (or their corresponding clusters) showed the most difference, while for grain yield ‘Nepal’ and ‘Southern Africa’ did. On the other hand, for days to flowering, the greatest difference was between ‘Nepal’ and ‘East Africa’.

Of the three last-mentioned traits, only plant height showed non-overlap between areas/clusters (for ‘East Africa’ vs. some other areas/clusters, where ‘East Africa’ accessions were, for example, on average 32% taller than accessions from ‘India’, **Fig. 4b**; the same non-overlap applied for the traits of leaf length, leaf number, leaf width [all greatest for ‘East Africa’] and number of productive tillers [fewest for ‘East Africa’], **Fig. S11**). In general, phenotypes were more differentiated between regions (Africa vs. Asia) than among geographic areas within regions, corresponding to genomic differentiation (**Fig. 2**) and consistent with combined phenotype centroids in **Fig. 4a**. Overall, each geographic area contained extensive phenotypic variation, with India containing the largest (**Tab. S7**). This is also be seen in the PCA and the boxplots (**Figs. 4** and **S11**).

### Genome-wide association analysis reveals a small number of significant associations and a range of candidate genes

Our GWAS detected 16 MTAs above the stringent Bonferroni threshold (-log_10_(0.05/8,778) = 5.24), 15 of which were chromosome-located. Twelve were associated with finger length (seven on chromosome 2B, three on 5B, one on 7B and one on 8B), two with days to flowering (one on 4B and one scaffold marker) and two with threshing percentage (one each on 4B and 6B). Manhattan and Q-Q plots for all 13 phenotypic traits are presented in **Fig. S12** and a list of all significant MTAs is given in **Tab. S8**. Most of the SNPs with significant MTAs that we identified by GWAS had a low MAF, in correspondence with our overall SNP panel (MAF < 0.2 for > 75% of all SNPs; see **Fig. S1d**). Therefore, care should be taken when interpreting our findings. The relatively small number of MTAs we detected could reflect the overall limited number of SNPs generated with the current genotyping strategy. All of the significant MTAs that we did detect were associated with the B sub-genome, even though it had a lower SNP density than the A subgenome (but the B sub-genome does have slower LD decay, see above). A list of 61 unique putative candidate genes revealed within a 25 kb interval both upstream and downstream of significant MTAs is given in **Tab. S8**, but we do not here explore these associations further.

### SNP effects clustering demonstrates different genetic architectures of phenotypic traits

Our exploration of SNP effects for phenotypic traits from RR-BLUP revealed two clusters of traits of different levels of complexity (**Fig. 5** and **Fig. S13**). Among traits with simpler genetic architecture were days to flowering and threshing percentage (consistent with the small number of MTAs detected for these two traits, see above). These are traits that do not follow a uniform distribution. Included among the traits with more complex genetic architecture (and of more uniform distribution) were finger length (which did still reveal MTAs in GWAS analysis) and grain yield (no MTAs in our analysis and known in other cereals to be highly polygenic; e.g., for wheat, see Brinton & Uauy, 2019).

**Fig. 5:**
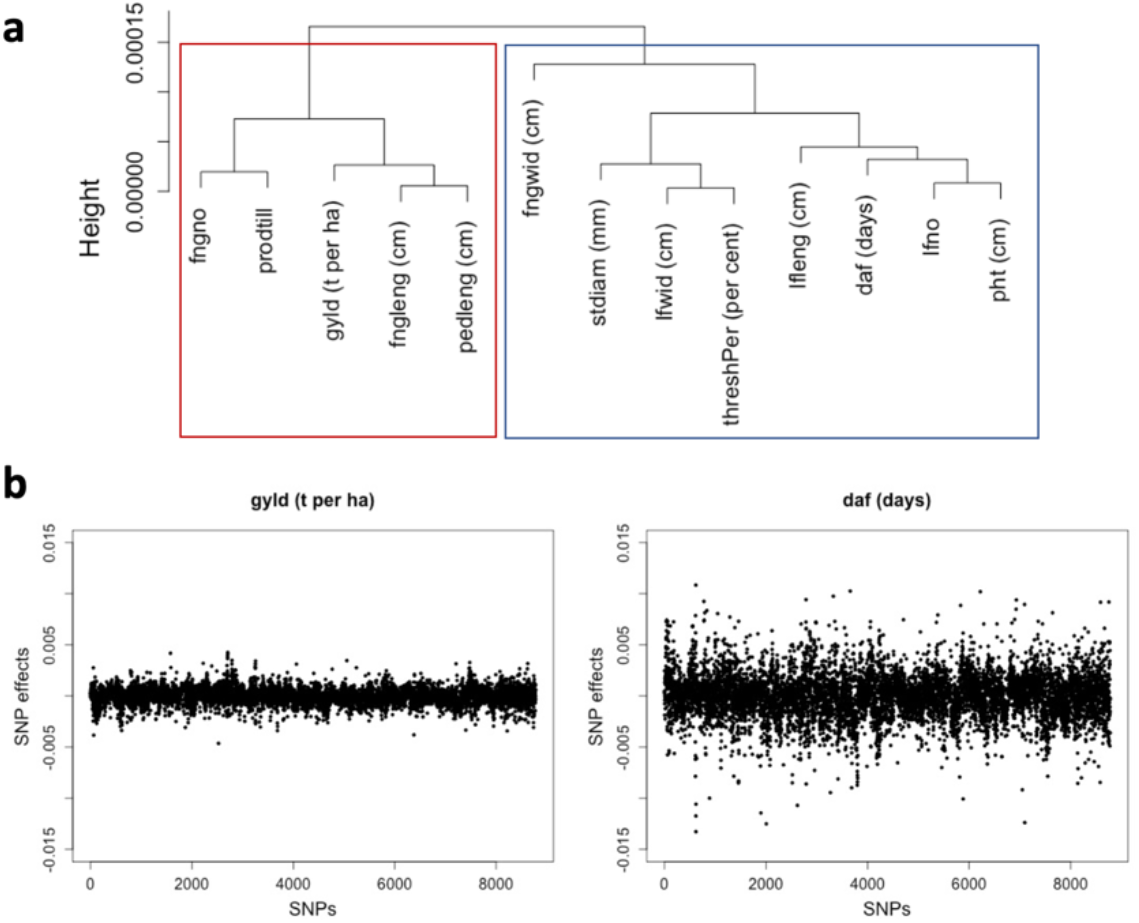
Clustering of SNP effects obtained from RR-BLUP to determine the complexity of agronomic traits. **a**, The UPGMA clustering algorithm was applied to the five statistical moments of SNP effects for all 13 agronomic traits. The cluster in red consists of traits with highly polygenic genetic architecture, and the cluster in blue consists of traits with simple-to moderately-complex genetic architecture. **b**, Examples of SNP effect distributions for the complex trait of gyld (grain yield) and the less complex trait daf (days to flowering). The results for the remaining 11 of the individual traits evaluated are shown in **Fig. S13**.

### Comparison of phenotypic and genome-wide gene diversity by geographic area supports varied post-domestication diversification pathways for finger millet

Our geographic-area-based comparison of phenotypic trait diversity with overall underlying gene diversity was based on the calculation of CV/H (i.e., standardized phenotypic variation per unit of genome-wide gene diversity; **Fig. 6**). The results showed that the ranking of values between geographic areas varied by trait, suggesting complex, context-specific diversification of the crop. The comparison of rankings of CV/H for finger number and plant height, for example, showed that the highest rank for the former was for ‘East Africa’ and for the latter was ‘Southern Africa’. Considering all 13 phenotypic traits, ‘East Africa’ most often of any geographic area ranked top for CV/H (in 6 cases) and ‘Nepal’ bottom (11 cases). This is consistent with the relatively low denominator (H) value for ‘East Africa’ compared to ‘Nepal’ (see above), and possibly indicates an ‘overexpression’ of phenotypic variation in the former region that is consistent with a longer domestication history. Days to flowering was the trait with the least spread in CV/H values for geographic areas, indicating that variation in this trait may be the best phenotypic proxy of underlying genomic diversity within geographic areas.

**Fig. 6:**
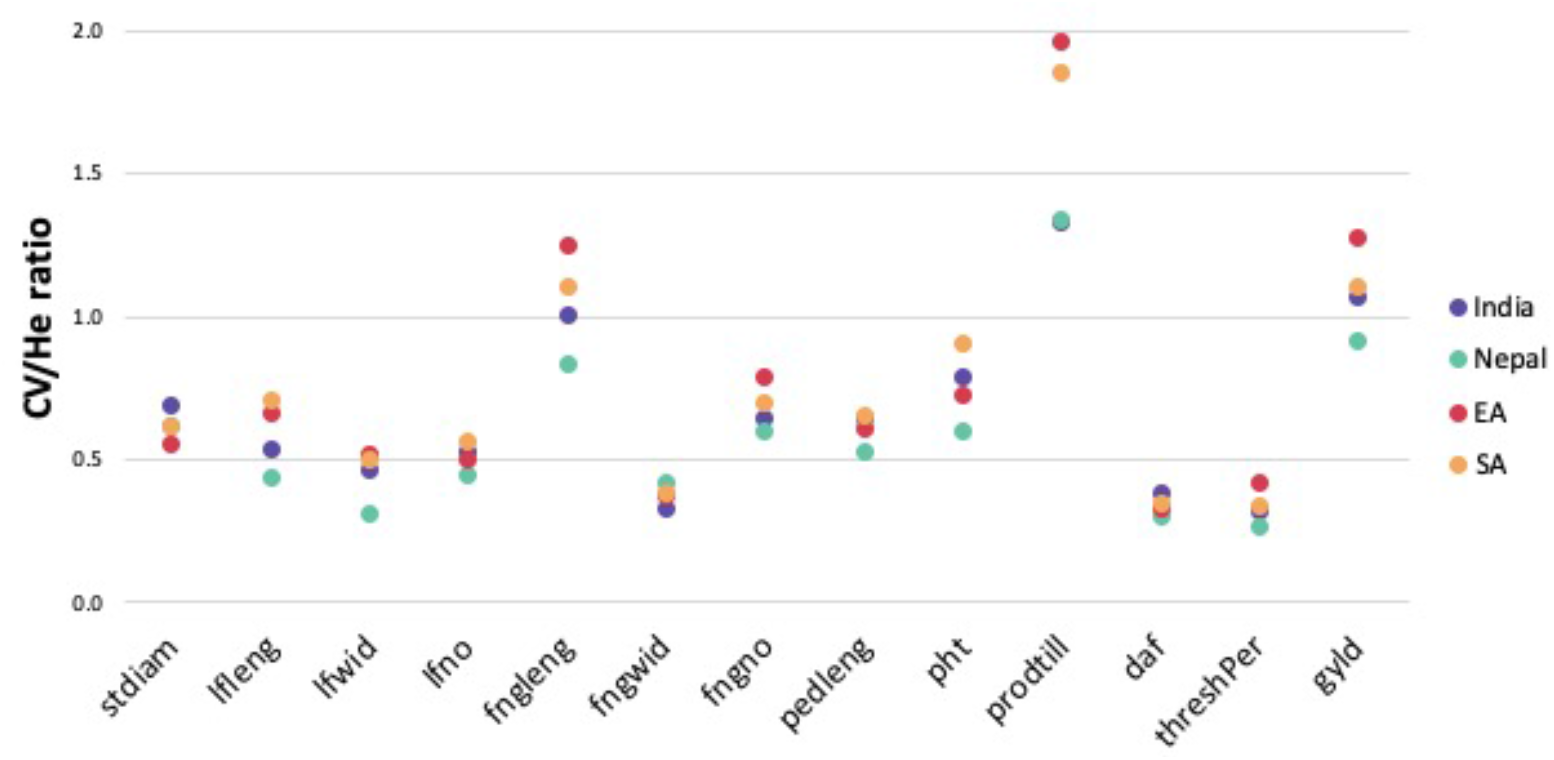
The ratio of the coefficient of phenotypic variance (CV) and genome-wide gene diversity (H) for 13 agronomic traits for four geographic areas. We used the ratios to inform us of finger millet’s selection history.

## DISCUSSION

Broadening crop production requires more focus on orphan crops. Orphan crops have an intrinsic value, but they can also provide broader lessons on crop domestication and crop diversification pathways (Dawson et al., 2019). Here, we have studied the orphan crop finger millet and have shown that a combination of modern and traditional methods can provide important insights relevant for the future development of the crop. Our approach also provides a model for work on other orphan crop species.

Past DNA-based studies of finger millet genetic diversity have generally been limited in scope, involving < 150 accessions and/or < 100 polymorphic loci. These studies include those of Dida et al. (2008), Arya et al. (2013), Manyasa et al. (2015), Ramakrishnan et al. (2016), Babu et al. (2018), Lule et al. (2018) and Pandian et al. (2018) who all applied simple sequence repeat (SSR) markers; and those of Gimode et al. (2016) and Sood et al. (2016) who used SSRs and SNPs. Despite their limited scope, these previous studies revealed a degree of locally- and regionally-structured genetic variation, including between Africa and Asia, and admixture between the two continents.

A restricted number of previous studies have also sought to explore marker-trait associations in finger millet. However, again the accessions and/or molecular markers involved were generally limited to small numbers, and studies did not have available a finger millet genome assembly to map markers to chromosome positions. These studies include those of Dida et al. (2021) who used a medium-sized panel of SNPs but only 52 East African accessions to search for blast disease resistance MTAs; Puranik et al. (2020) who used a large panel of SNPs but assessed only 190 genotypes to search for grain nutrient-content-related MTAs; Sharma et al. (2018) who applied a medium-sized panel of SNPs to only 113 accessions to search for MTAs in 14 agro-morphological traits; and Tiwari et al. (2020) who, beginning with the same SNPs and germplasm panel of Sharma et al. (2018), explored marker associations with grain protein content. Despite their limited scope, these previous studies indicated some MTAs. These were identified under varying degrees of stringency and certainty levels, applying for example candidate gene sequence homologies for characterization.

Our analysis was based on both an extensive germplasm panel and a broad genome-wide (and sub-genome-located) SNP marker set. This was combined with extensive field phenotypic assessment, the use of a finger millet genome assembly, and the application of a suite of population genomic tools. Overall, this provides us with a more complete picture than has been possible previously of genetic variation in the finger millet crop. Our analysis supports previous studies that have identified geographically-structured variation. However, it also sheds more light on the apparently complex evolutionary history of finger millet and its varied within and between region diversification pathways, thereby providing new information to support future breeding. Below, we focus further discussion on two aspects of particular interest that expand our current understanding of finger millet: i) polyploidy and sub-genome-specific diversification; and ii) geographically-specific relationships between phenotype and genotype embracing chromosome-, sub-genome-, and genome-level diversity, with a range of individual phenotypic traits and combinations of traits.

### Sub-genome-specific diversification and polyploidy

Hybridization between different genomes, followed by chromosome doubling to generate polyploids, has played a crucial role in cereal crop evolution (Levy & Feldman, 2002). This has been well studied for major cereals such as allohexaploid wheat (Feldman & Levy, 2012). However, the full origin of the finger millet allotetraploid genome is not well understood (Zhang et al., 2019). The allopolyploidization event from diploid A and B sub-genome progenitors to form the tetraploid crop ancestor (*E. corocana* subsp. *africana*) is believed to have occurred in East Africa (de Wet et al., 1984). Traditionally, such polyploidization events are thought to be rare in plant’s histories and initially lead to genomic bottlenecks in the derived organisms (Stebbins, 1950). In the current study, we could not compare relative diversity levels of the cultivated finger millet with the descendants of its wild tetraploid progenitor or previous diploid progenitors. However, previous studies found genomic diversity in cultivated finger millet to be significantly lower than in wild *E. coracana* subsp. *africana* (e.g., Gimode et al., 2016). We were nevertheless able to explore for the first time in finger millet the sub-genomic and chromosome-specific diversity patterns in the crop by geographic area of sampling, which may inform on its evolution. We detected the highest level of genetic diversity in the B sub-genome overall in ‘ Southern Africa’, with exceptionally high diversity levels along two chromosomes (5B and 9B). In contrast, for the A sub-genome, ‘ Southern Africa’ only ranked third in diversity, while diversity for ‘East Africa’ ranked lowest for both sub-genomes.

Our new observations are consistent with broadening of diversity in specific parts of the finger millet genome as the crop developed new adaptations during expansion from East Africa to new areas in Africa and in Asia. This may have been facilitated by the polyploidization process that enables mechanisms such as inter-genomic transfer through translocation, recombination, and transposition to support rapid evolution (as outlined by Levy & Feldman, 2002). Our results could also indicate a secondary contact in the ‘Southern Africa’ region between the crop and its immediate progenitor, or perhaps new introgressions from other co-located *Eleusine* species that have specifically targeted the B sub-genome and could have been aided by polyploidization. Both wild *E. coracana* subsp. *africana* (i.e., the considered most likely immediate progenitor) and a range of other *Eleusine* species are sympatric with the finger millet crop in southern Africa (**Fig. 1**), providing opportunities for introgression. This type of process has been observed in other cereals, including wheat (He et al., 2019), and would be consistent with genotyping-by-sequencing in finger millet that has revealed chromosomal rearrangements between sub-genomes (Qi et al., 2018). Regardless of the cause, the patterns of sub-genome-specific diversity observed in our study suggest complexity in finer millet’s evolution. Genomic exploration of *Eleusine* germplasm panels containing extant descendants of the known and putative crop wild progenitors may further elucidate this complex demographic history.

### Region-specific genomic-phenotypic relationships

Our analysis shows complex relationships between genomic and phenotypic variation in finger millet across the four geographic areas we sampled. This is consistent with a crop that has been subject to millennia of domestication and has experienced different selection pressures based on particular human preferences and production environments in different locations. Complex relationships between genetic variation and phenotypic variation are common in crops (e.g., Kozak et al., 2011), and our results were therefore not unexpected.

Our analysis of genetic diversity along sub-genomic chromosome homeologs indicated differences in relative diversity for particular geographic areas. The subsequent pooling of chromosome diversity at the sub-genome level also revealed different diversity rankings by location. When levels of geographic-area-based phenotypic variation were compared with genome-wide gene diversity (through the calculation of CV/H), the results suggested context-, trait-, and trait-combination-specific diversification pathways for the crop. In addition, the ‘East Africa’ area most often ranked top for CV/H values, consistent with a longer cultivation history here than elsewhere. The profiles of phenotypic traits alone that varied between geographic areas by individual trait further suggest multiple selection pressures both between and within areas, with the broad variation often captured within areas indicating local differential selection.

Finally, patterns of differentiation between geographic areas also varied for genomic data compared to phenotypic data, further supporting complex demographic processes. For example, genome-wide SNPs revealed the highest differentiation between ‘Southern Africa’ and ‘Nepal’ accessions, but combined phenotypic data indicated the greatest level of difference between ‘India’ and ‘East Africa’. This perhaps illustrates that these last two areas are the strongest foci for context-specific crop development and/or are most agro-ecologically differentiated.

### Concluding remarks

This study has revealed distinct genomic and phenotypic variation patterns in finger millet that suggest complex diversification pathways for the crop. Our data provide a firmer basis for the future development of finger millet. The information can, for example, help guide the determination of heterotic groups to produce potentially hybrid finger millet varieties in the future (Labroo et al., 2021; Mackay et al., 2021). Hybridization between ‘East Africa’ and ‘Nepal’ or ‘Southern Africa’ accessions may also generate useful genetic variation for use in the genetically narrow East Africa region. A study of *Eleusine* germplasm panels containing known and putative crop wild progenitors is required to understand the crop’s demographic history further. In our ongoing research, we seek to understand the nature and extent of genotype-by-environment interactions in finger millet, to guide whether more local germplasm panels would be better suited for GWAS, and to assess the prospects for climate change adaption of the crop. As genomic resources continue to develop for finger millet, breeding material selection approaches that assign genomic estimated breeding values to each separate sub-genome also open a possibility for weighted sub-genome selection (Santantonio et al., 2019).

## Abbreviations

AMOVA: analysis of molecular variance;
CV: coefficient of variation;
DAPC: Discriminant Analysis of Principal Components;
DArTseq: Diversity Arrays Technology sequencing;
GRM: genomic relationship matrix;
GWAS: genome-wide association study;
LD: linkage disequilibrium;
MAF: minor allele frequency;
MLM: mixed linear model;
MTA: markertrait association;
NJ: neighbour-joining;
PCA: principal component analysis;
RR-BLUP: ridge regression-best linear unbiased genomic prediction;
SNP: single nucleotide polymorphism

## ACKNOWLEDGEMENTS

The authors are thankful for helpful comments from Ian Mackay and Rajiv Sharma, and technical assistance from Ann Murithi and Erick Owuor Mikwa. We also thank Peter Bradbury for adding a bespoke filtering option in TASSEL for removing SNPs with a heterozygosity level greater than 2pq × (1 – F).

## AUTHOR CONTRIBUTIONS

JB, IKD and DAO conceptualized the study. HFO coordinated the collection of field phenotype data. DAO coordinated the generation and initial screening of molecular marker data. SMJ carried out genomic DNA extractions of the 458 samples. JB conducted further data screening and all statistical analyses. IKD, RCG, JBu, GG and DAO contributed to determining appropriate analysis methods and interpreting findings. JB and IKD wrote the first draft of the paper that was then contributed to by all other authors.

## SUPPLEMENTAL MATERIAL

**Supplemental File S1** contains supplemental tables (**Tables S2,5,6**) and figures (**Figs. S1-13**).

**Supplemental File S2** contains supplemental **Tables S1,3,4,7,8**.

## DATA AVAILABILITY

All data generated and analyzed during this study are included in this published article and its supplemental material. The raw phenotypic and DArTseq SNP data were deposited into Dryad Digital Repository (http://datadryad.org/).

