## SupplementalFileS1 for "Genomic and phenotypic characterization of finger millet indicates a complex diversification history"

**Supplemental Figure 1. Quality control of genomic data.** Measures were taken in the order presented here. **a,** Removal of accessions with missingness rate > 30%, **b**, Removal of SNPs with missingness rate > 30%, **c**, **d,** Filtering for minor allele frequency < 0.01, before (**c**) and after (**d**) application, and **e**, **f,** Filtering for SNPs with a heterozygosity level greater than 2pq × (1-F) (red curve), before (**e**) and after (**f**) application. Vertical lines on plots **a** to **d** represent the cut-off used in filtering, while in **e** and **f** this is represented by the red curve. The green curve represents the Hardy-Weinberg Equilibrium assumption.

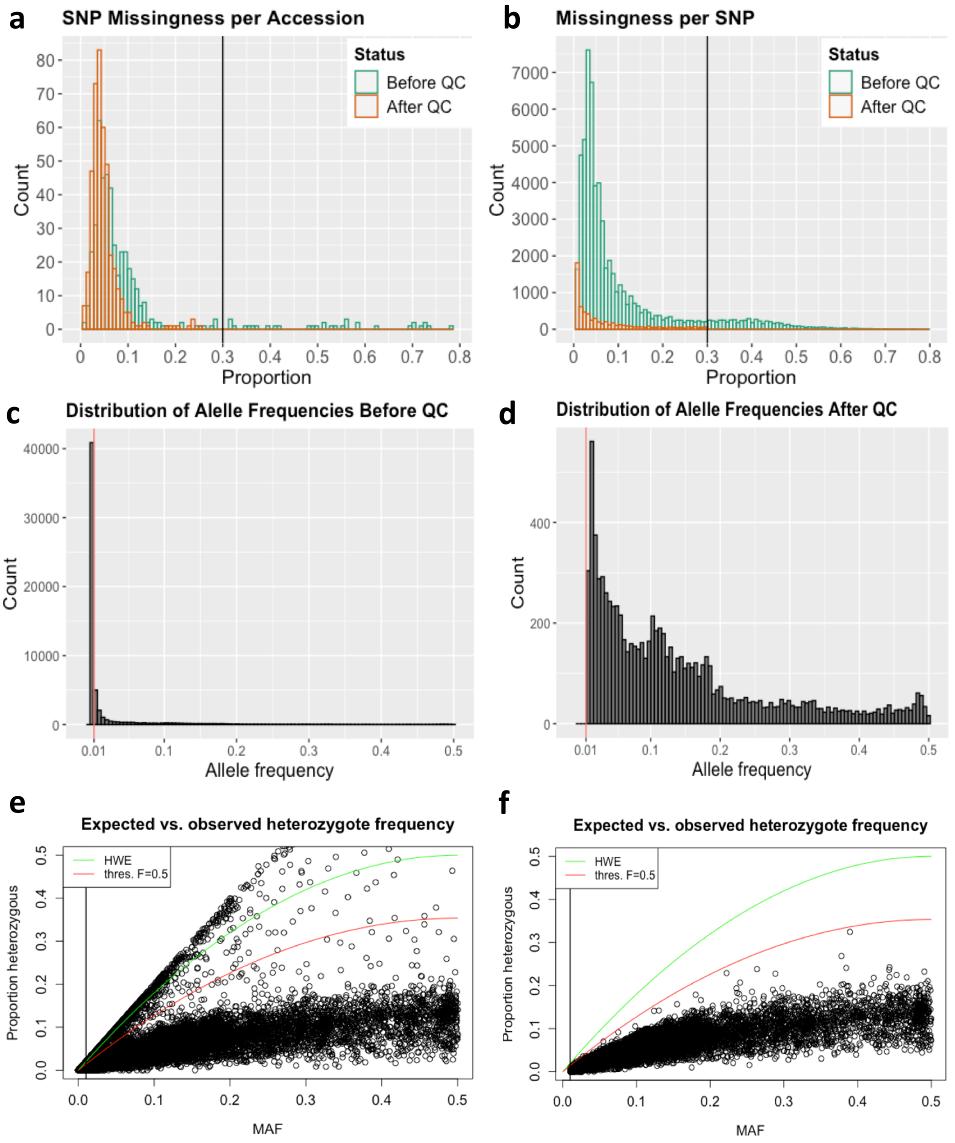

**Supplemental Figure 2. Scree plot of the first ten principal components from a PCA analysis of genetic structure.** A bend (indicated by the vertical line) appeared at PC = 4. This was considered the optimal number to account for genetic structure in subsequent analyses (i.e. DAPC and GWAS).

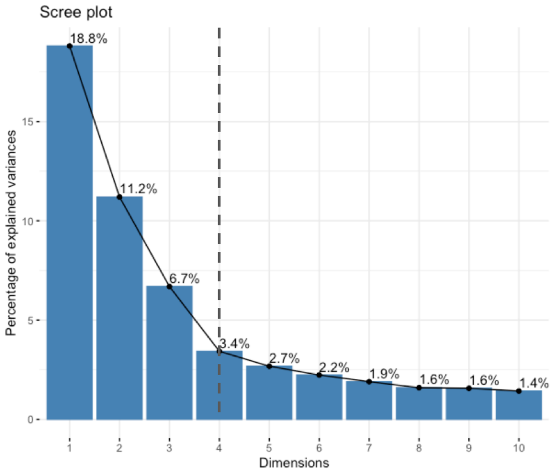

**Supplemental Figure 3. Plot of Bayesian Information Criterion (BIC) values for *k*-value tested in the discriminant analysis of DAPC.** These were generated by “find.clusters” in the R package “adegenet” (Jombart, 2008). A bend (indicated by the vertical line) appeared at $k = 4$. This was considered to be the optimal *k*-value.

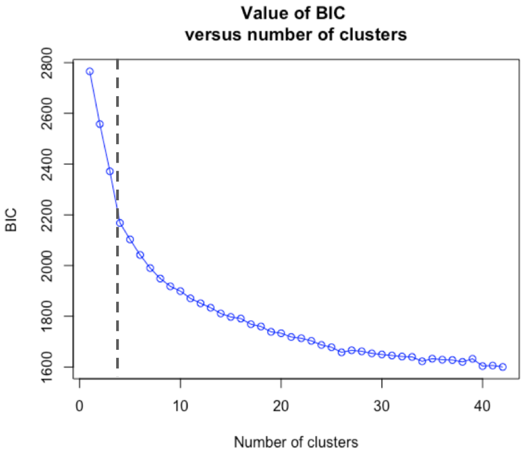

**Supplemental Table 2. Description of 13 agronomic traits measured for 423 finger millet accessions.**

| Trait | Abbrev. | Type | Method of collection |
|---|---|---|---|
| Stem diameter (mm) | stdiam | Morph. | Measured across centre between third and fourth node on the main tiller at dough stage |
| Leaf length (cm) | lfleng | Morph. | Measured from ligule to tip of the fourth leaf blade on the main tiller at head emergence stage |
| Leaf width (cm) | lfwid | Morph. | Measured across centre of the fourth leaf blade on the main tiller at head emergence stage |
| Leaf number | lfno | Morph. | The number of leaves on main tiller at dough stage |
| Finger length (cm) | fngleng | Morph. | Finger length is measured from the base of the longest finger on the main tiller to the tip at dough stage |
| Finger width (cm) | fngwid | Morph. | Measured across centre of the longest finger on the main tiller at dough stage |
| Finger number | fngno | Morph. | The number of mature fingers on main ear at dough stage |
| Peduncle length (cm) | pedleng | Morph. | From top most node to base of the thumb finger at dough stage |
| Plant height (cm) | pht | Morph. | Measured from ground level to tip of the inflorescence of the main tiller at dough stage |
| Productive tillers | prodtill | Phys. | Number of basal tillers which bear mature ears at dough stage |
| Days to flowering* | daf | Phys. | From sowing to stage when ears have emerged from 50% of main tillers |
| Threshing percentage (%)* | threshPer | Yield | Calculated as dry panicle weight [g] divided by grain yield per plot [g] after harvesting |
| Grain yield (t/ha)* | gyld | Yield | Weight of total grain yield of five tagged plants was recorded and yield per hectare was calculated after harvesting stage |

* measurements were taken on the plot basis and not based on the average of 5 random plants in the plot; *Morph.*, morphological trait; *Phys.*, physiological trait; *Yield*, yield-related trait

**Supplemental Figure 4. Distribution of DArTseq generated SNPs on finger millet chromosomes.** The figure includes 8,096 SNPs (excluding 682 scaffold SNPs) which remained after quality control. The colour key shows the density of SNPs per Mb.

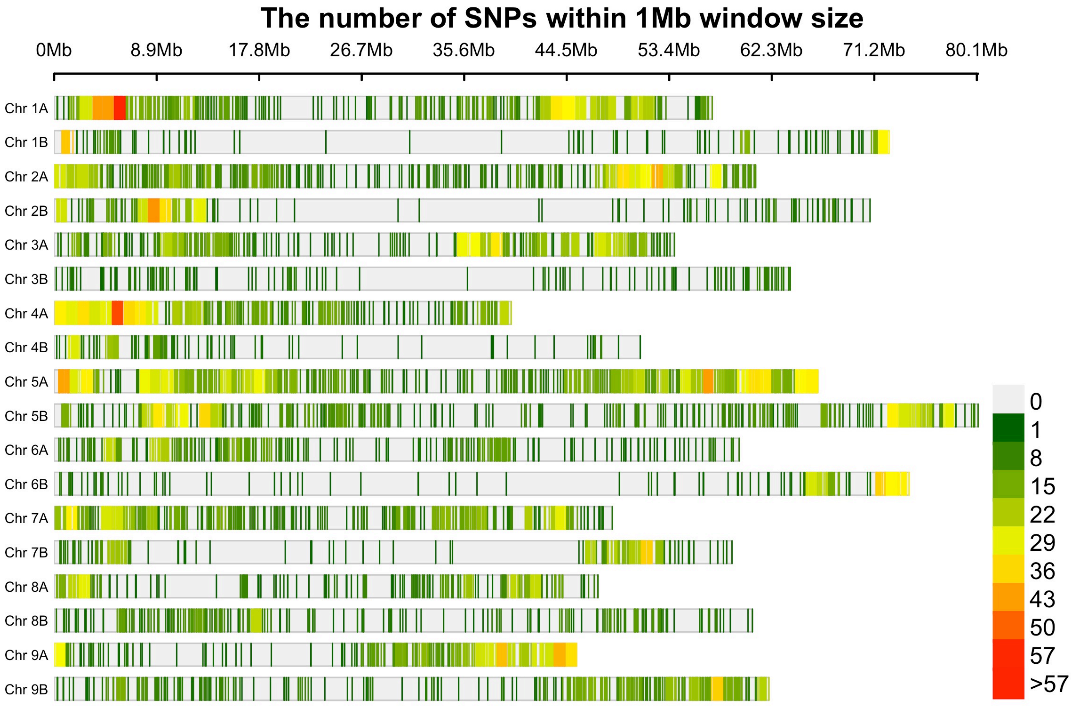

**Supplemental Figure 5. Linkage disequilibrium decay in finger millet**. Plots are either uncorrected (left side figures) or corrected (right side figures) for relatedness. **a**, Genome-wide values. **b**, Sub-genomes values. **c**, Single chromosome values. Values for all possible pairwise combinations of SNPs were plotted against genetic distance in megabases (Mb). The dotted horizontal line shows the $r^2$ threshold (0.2), and the vertical line the critical value at which the decay curve (red) intercepts the threshold, as indicated by the number on each plot.

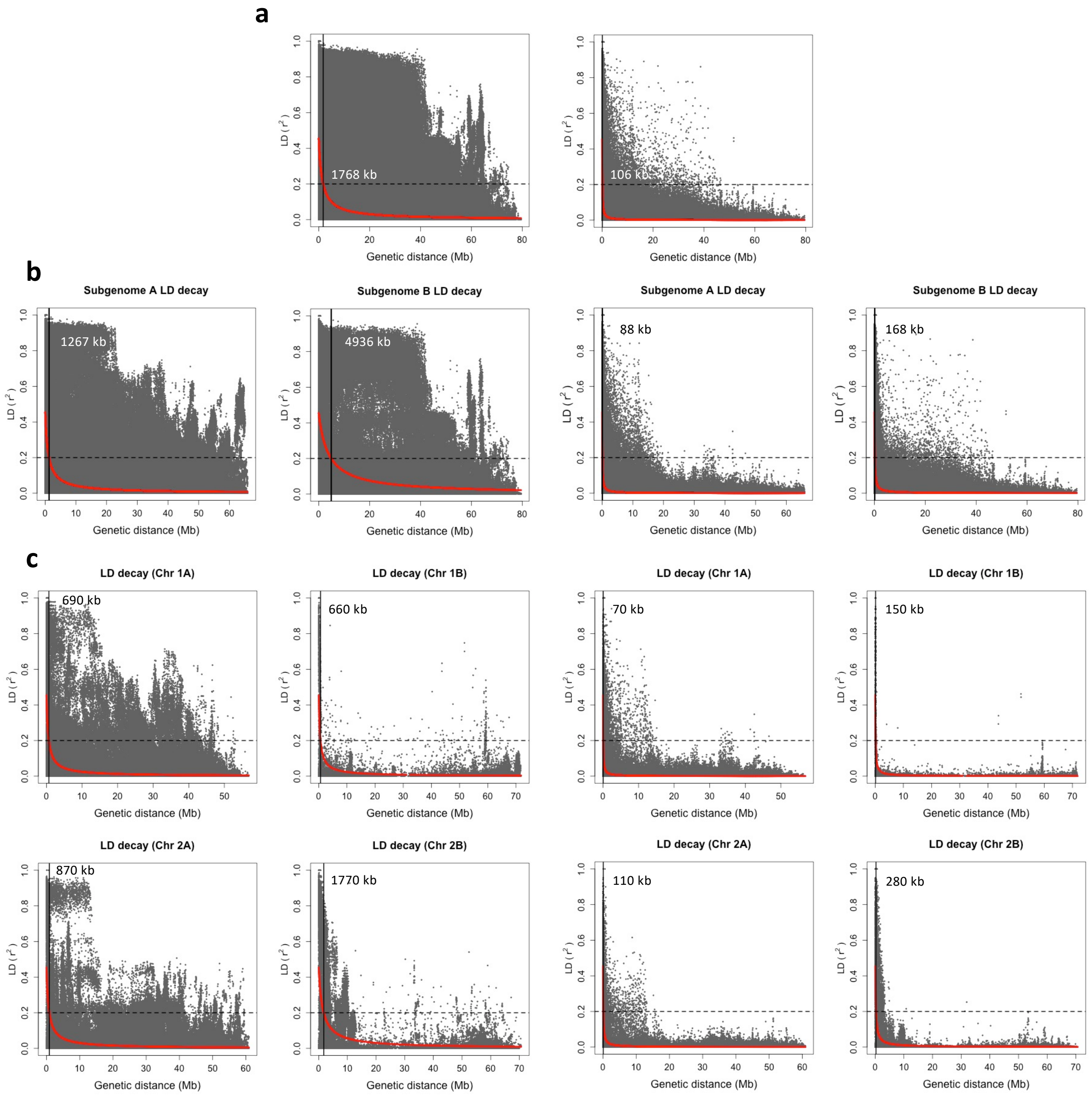

**Supplemental Figure 5.** Continued

c

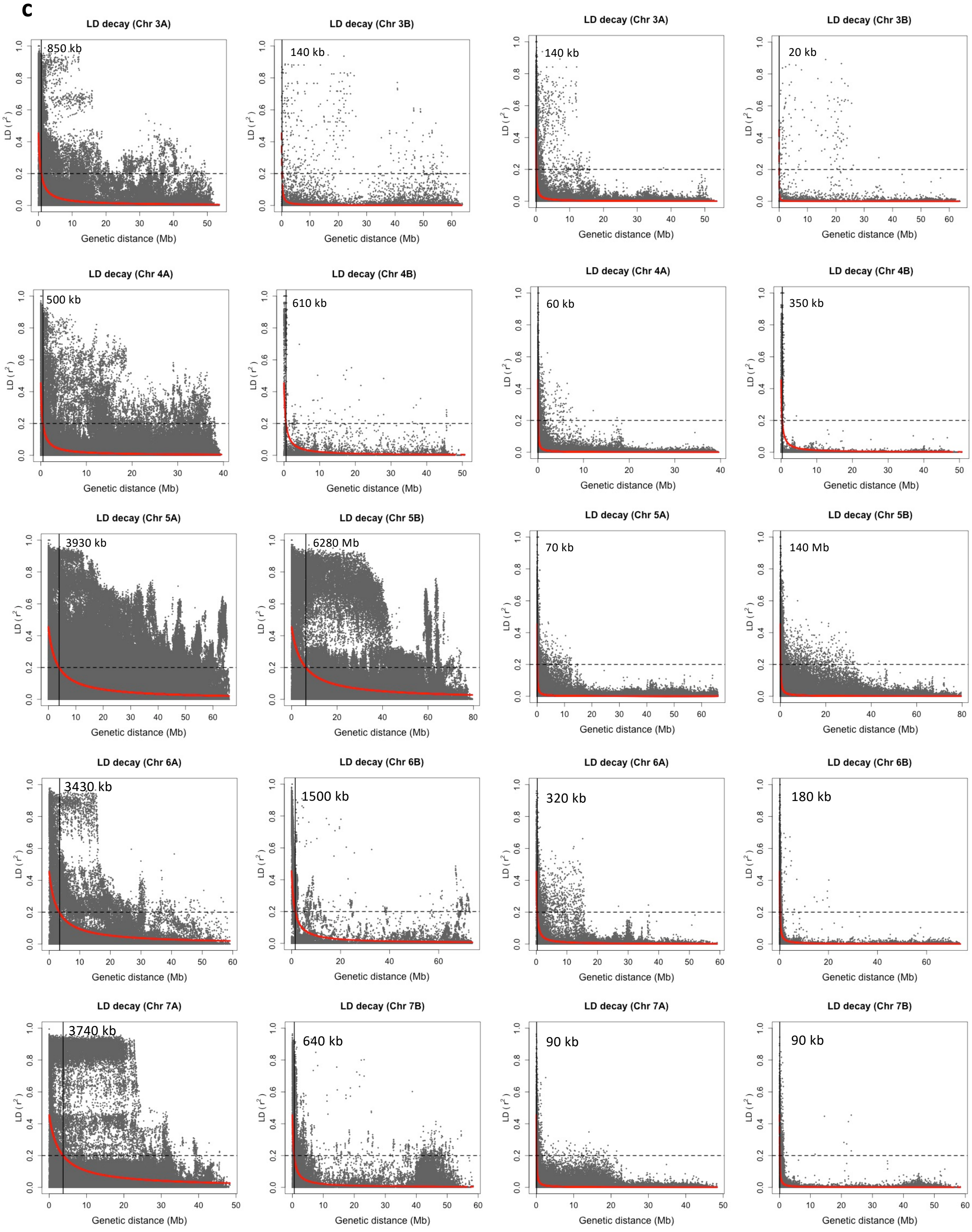

**Supplemental Figure 5.** Continued

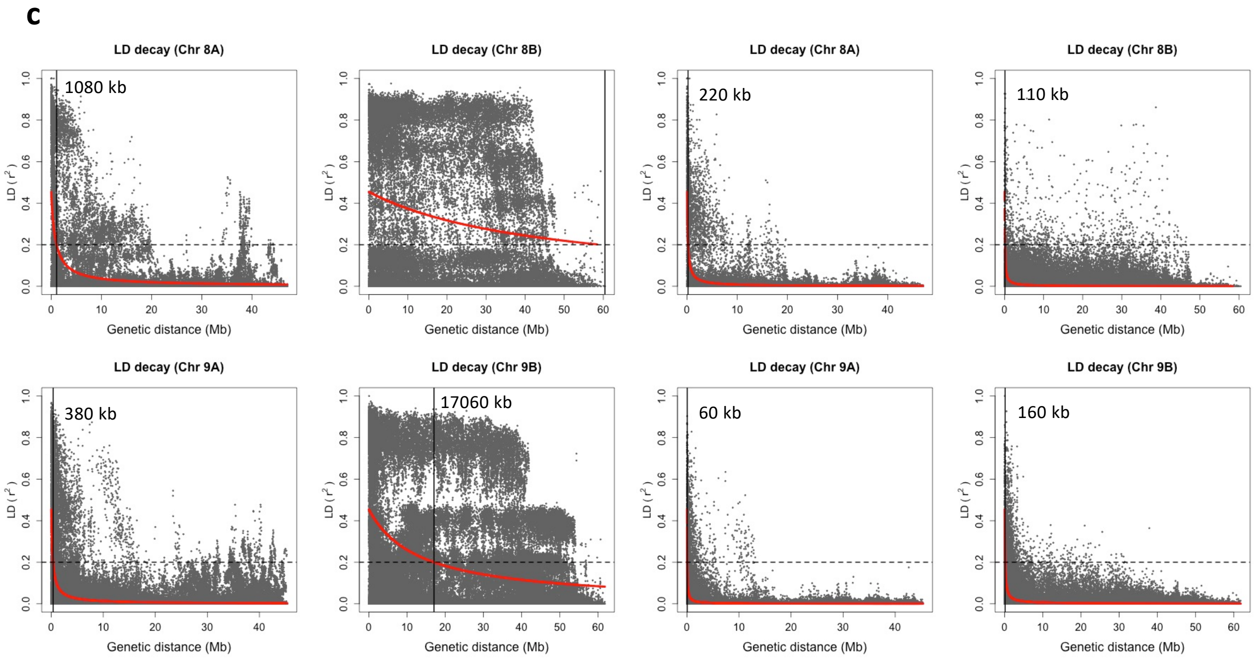

**Supplemental Figure 6. Heatmap and dendrogram of the genomic relationship matrix.** The matrix was estimated based on 8,778 SNPs and 423 finger millet accessions. Both rows and columns represent accessions. UPGMA clustering was applied to generate dendrograms. The coloured horizontal bar represents to which geographic area each accession belongs, with overall groups represented by squares in the matrix. The histogram represents the distribution of values in GRM.

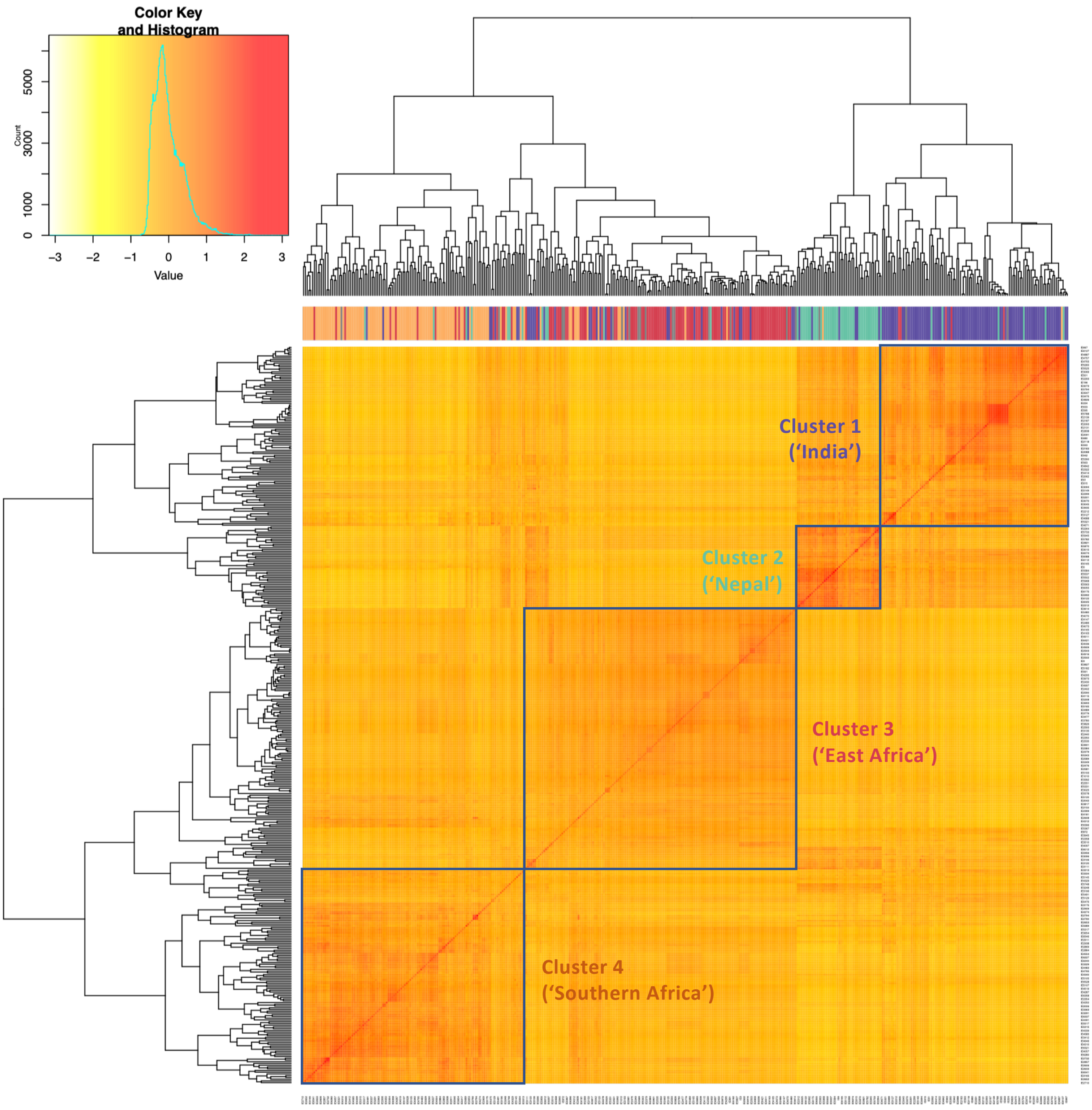

**Supplemental Figure 7. Phylogenetic neighbour-joining trees for the two finger millet sub-genomes. a**, Sub-genome A; **b**, Sub-genome B. Tip colour coding represent the geographic area to which each accession belongs.

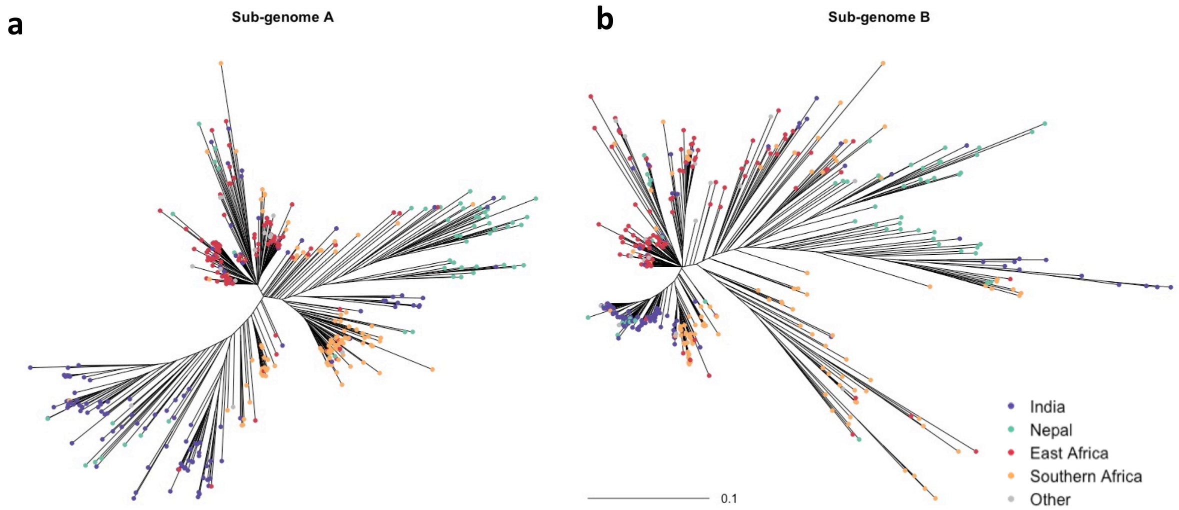

**Supplemental Figure 8. Visualization of the matrix of residuals from Treemix analysis. a**, Residual fit from the maximum likelihood tree without migration events fitted. **b**, with one migration event fitted.

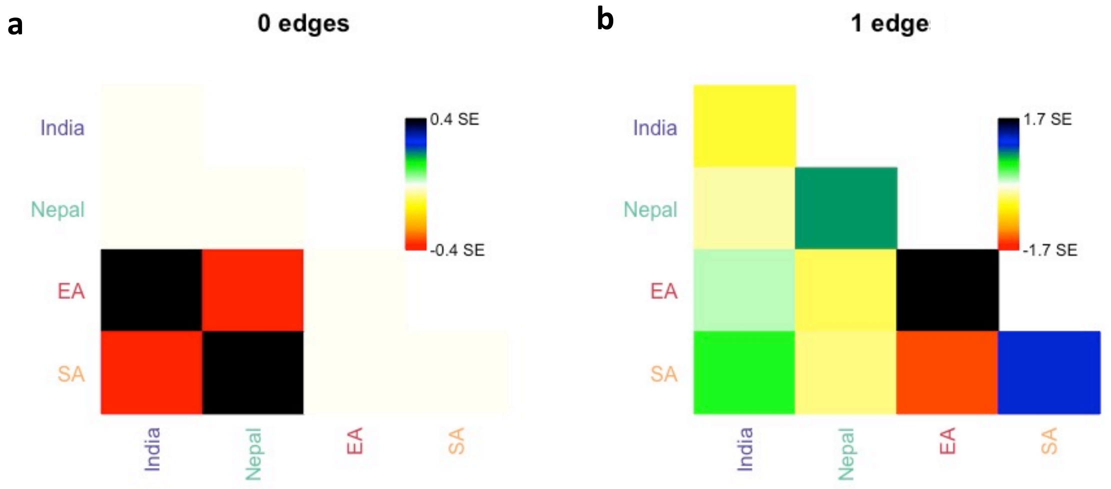

**Supplemental Figure 9. Genome-wide scans of diversity along finger millet chromosomes for geographic areas. a**, Nucleotide diversity ($\pi$) for each geographic area: ‘Nepal’ (turquoise line), ‘India’ (purple line),’ East Africa’ (red line) and ‘Southern Africa’ (orange line). **b**, Differentiation ($F_{ST}$) is given for each pairwise comparison between geographic areas. Both $\pi$ and $F_{ST}$ estimates were calculated using 1 Mb non-overlapping windows in VCFtools. **c**, The number of SNPs per window along the genome.

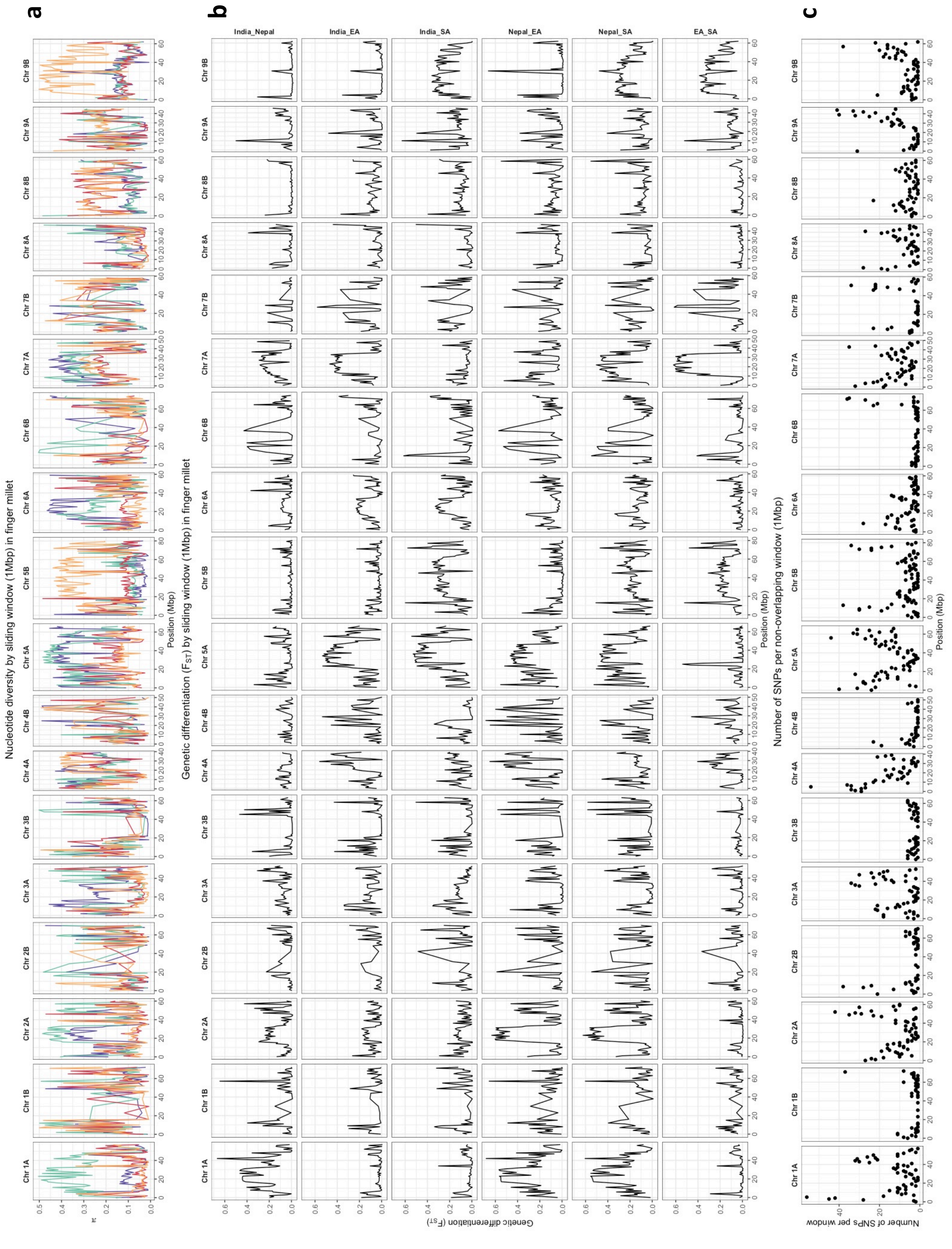

**Supplemental Table 5. Pairwise genome-wide $F_{ST}$ values for finger millet accessions sampled from geographic areas.**

| Region | India | Nepal | EA | SA |
|---|---|---|---|---|
| **India** | – | 0.069 | 0.104 | 0.131 |
| **Nepal** | – | – | 0.137 | 0.167 |
| **EA** | – | – | – | 0.087 |
| **SA** | – | – | – | – |

**Supplemental Table 6. Summary statistics of best linear unbiased estimates (BLUEs) for 13 agronomic traits in finger millet.**

| Trait | Trait unit | Descriptive statistics | | | | Variance parameters | | | |
|---|---|---|---|---|---|---|---|---|---|
| | | Mean | SD | Min | Max | CV | $\sigma^2_G$ | $\sigma^2_E$ | $H^2$ |
| stdiam | cm | 8.41 | 1.32 | 4.13 | 12.66 | 15.72 | 1.14 | 0.98 | 0.67 (0.03) |
| lfleng | cm | 41.95 | 7.53 | 22.23 | 63.99 | 17.95 | 46.40 | 13.90 | 0.85 (0.01) |
| lfwid | cm | 1.21 | 0.15 | 0.74 | 1.66 | 12.80 | 0.02 | 0.01 | 0.80 (0.02) |
| lfno | number | 10.78 | 1.55 | 3.60 | 16.81 | 14.40 | 1.46 | 1.44 | 0.63 (0.03) |
| fngleng | cm | 5.40 | 1.51 | 2.82 | 14.89 | 27.94 | 2.12 | 0.29 | 0.94 (0.01) |
| fngwid | cm | 0.93 | 0.09 | 0.57 | 1.18 | 9.63 | 0.00 | 0.01 | 0.62 (0.03) |
| fngno | number | 7.07 | 1.26 | 3.67 | 11.87 | 17.89 | 1.09 | 0.86 | 0.68 (0.03) |
| pedleng | cm | 24.21 | 4.05 | 10.17 | 37.41 | 16.74 | 12.92 | 5.49 | 0.81 (0.02) |
| pht | cm | 74.60 | 17.32 | 24.04 | 123.84 | 23.22 | 269.56 | 45.13 | 0.92 (0.01) |
| prodtill | cm | 6.61 | 3.19 | 2.07 | 18.64 | 48.25 | 6.83 | 5.18 | 0.70 (0.03) |
| daf | days | 69.47 | 6.05 | 45.84 | 92.20 | 8.71 | 34.65 | 3.15 | 0.95 (0.00) |
| threshPer | % | 75.88 | 6.51 | 47.27 | 89.65 | 8.58 | 16.06 | 47.78 | 0.36 (0.05) |
| gyld | ton/ha | 3.36 | 0.95 | 0.76 | 6.02 | 28.33 | 0.34 | 1.01 | 0.35 (0.05) |

*SD* standard deviation, *Min* minimum, *Max* maximum, *CV* coefficient of variance, $\sigma^2_G$ genetic variance, $\sigma^2_E$ residual variance, $H^2$ broad sense heritability, *stdiam* stem diameter, *lfleng* leaf length, *lfwid*, leaf width, *lfno* leaf number, *fngleng* finger length, *fngwid* finger width, *fngno* finger number, *pedleng* peduncle length, *pht* plant height, *prodtill* production tillers, *daf* days to flowering, *thershPer* threshing percentage, *gyld* grain yield

**Supplemental Figure 10. Distribution of phenotypic variation and correlations for 13 agronomic traits in finger millet.** Profiles are based on best linear unbiased estimates (BLUEs) of traits. The distribution of each trait is shown on the diagonal, the bivariate scatter plots with a fitted line are displayed beneath the diagonal, and the value of the correlation between two traits with its significance level (* $p < 0.05$, ** $p < 0.01$, and *** $p < 0.001$) above the diagonal. *stdiam,* stem diameter; *lfleng,* leaf length; *lfwid*, leaf width; *lfno,* leaf number; *fngleng,* finger length; *fngwid,* finger width; *fngno,* finger number; *pedleng,* peduncle length; *pht,* plant height; *prodtill,* production tillers; *daf,* days to flowering; *thershPer,* threshing percentage; *gyld,* grain yield.

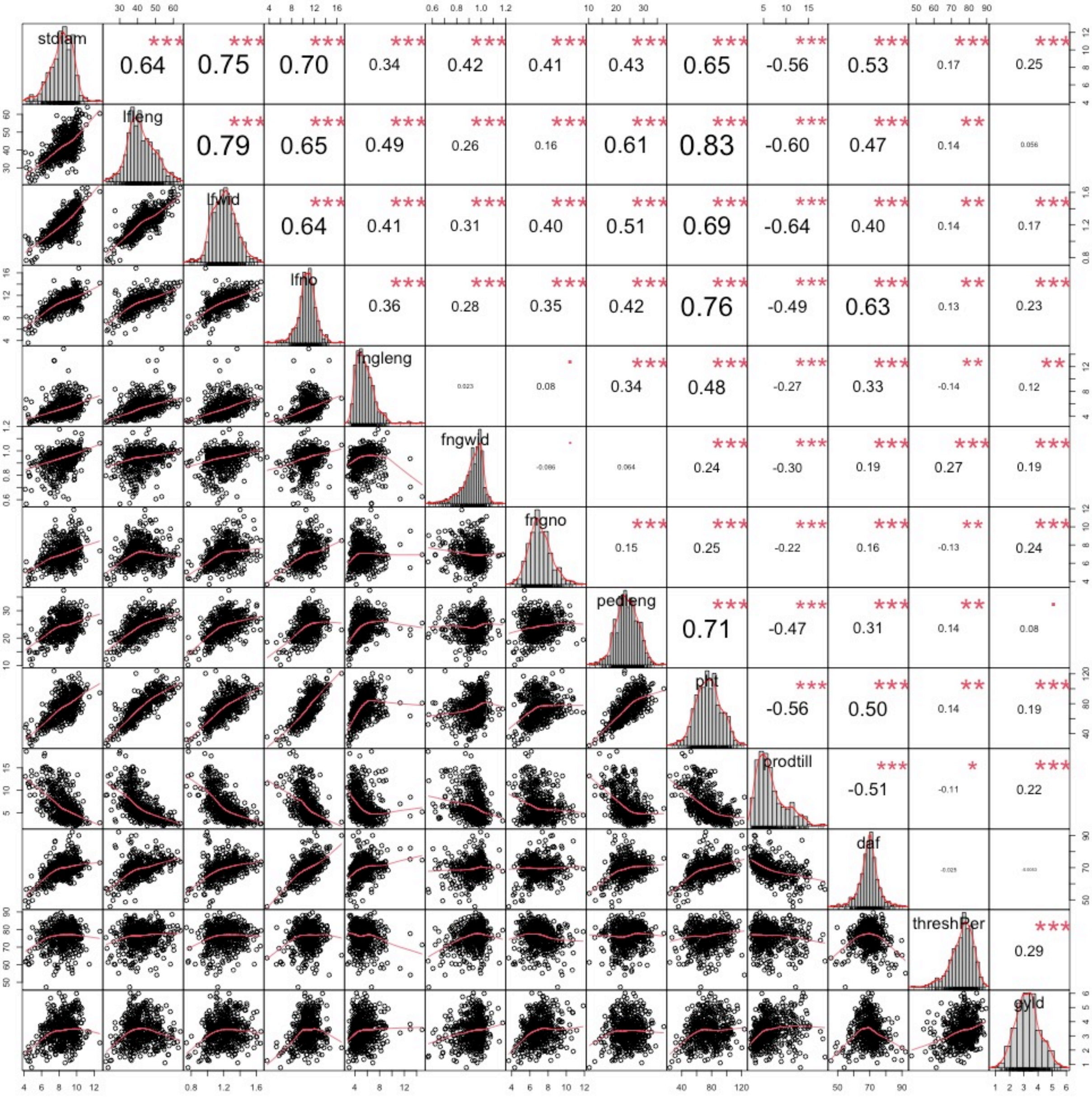

**Supplemental Figure 11. The statistical distribution of 10 agronomic traits measured in 423 finger millet accessions.** Boxplots are shown according to genetic clusters identified by discriminant analysis (purple boxplots), geographic areas of origin (orange boxplots) and total phenotypic variation (green boxplots). A bar and a point inside each boxplot represent median and mean, respectively. The remaining three traits to total 13 in all are presented in **Fig. 4**. *stdiam,* stem diameter; *lfleng,* leaf length; *lfwid*, leaf width; *lfno,* leaf number; *fngleng,* finger length; *fngwid,* finger width; *fngno,* finger number; *pedleng,* peduncle length; *prodtill,* production tillers; *thershPer,* threshing percentage.

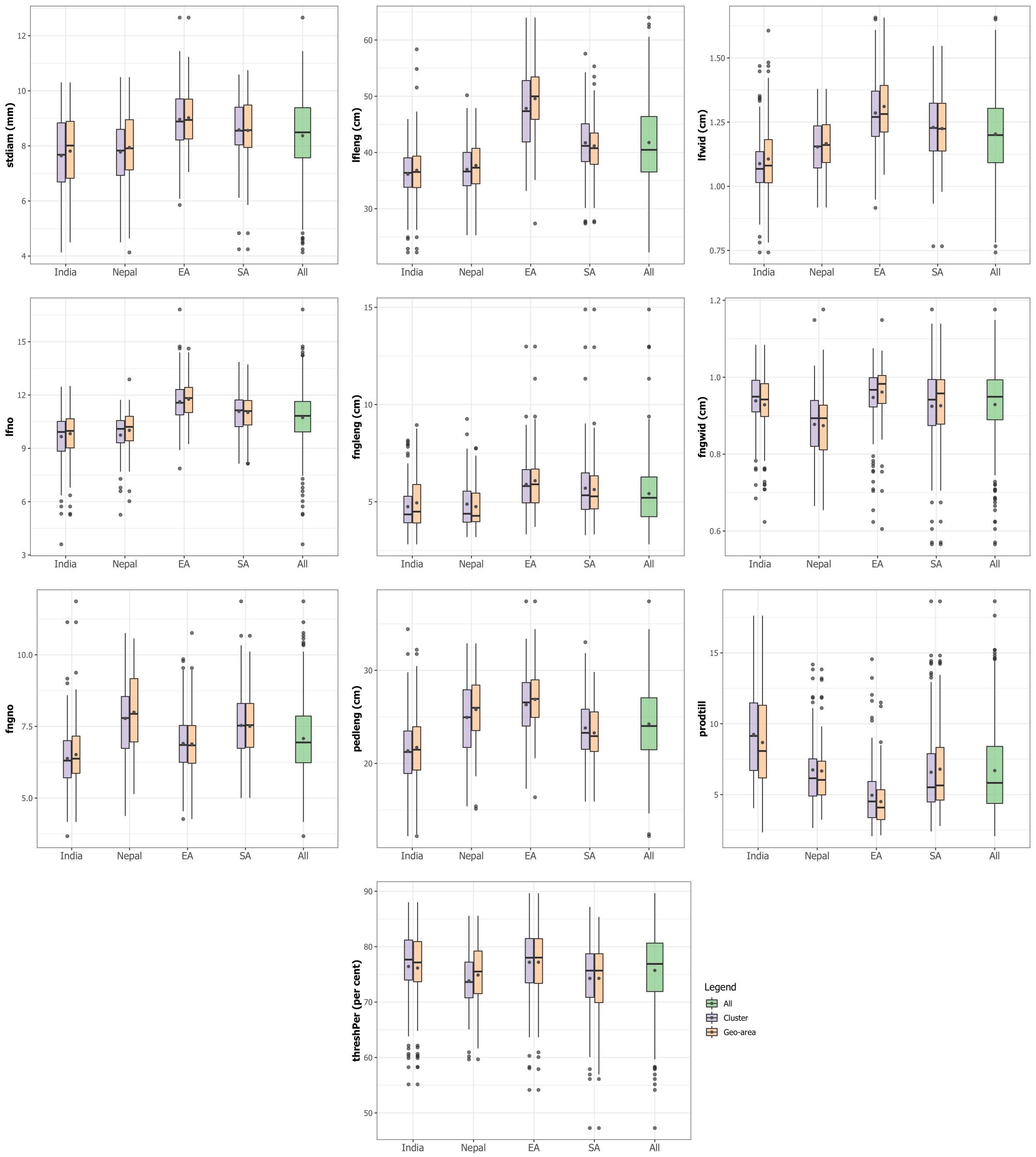

**Supplemental Figure 12. GWAS results presented with Manhattan plots and quantile-quantile plots for 13 agronomic traits**. The red horizontal line on Manhattan plots (left side plots) indicates a Bonferroni threshold of 5% significance. On quantile-quantile plots (right side plots, the red diagonal line represents the theoretical reference distribution. *stdiam,* stem diameter; *lfleng,* leaf length; *lfwid*, leaf width; *lfno,* leaf number; *fngleng,* finger length; *fngwid,* finger width; *fngno,* finger number; *pedleng,* peduncle length; *pht,* plant height; *prodtill,* production tillers; *daf,* days to flowering; *thershPer,* threshing percentage; *gyld,* grain yield.

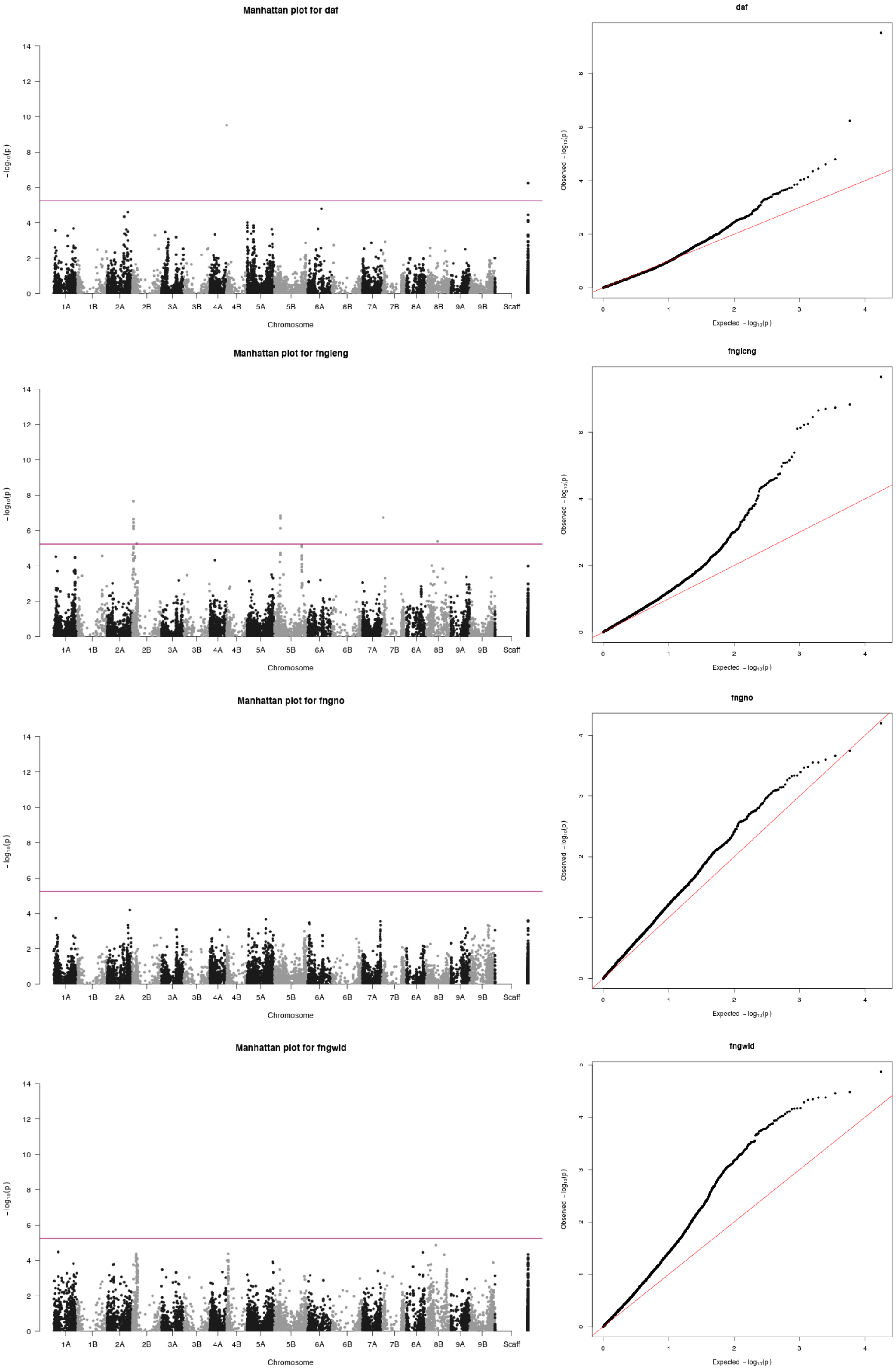

**Supplemental Figure 12.** Continued

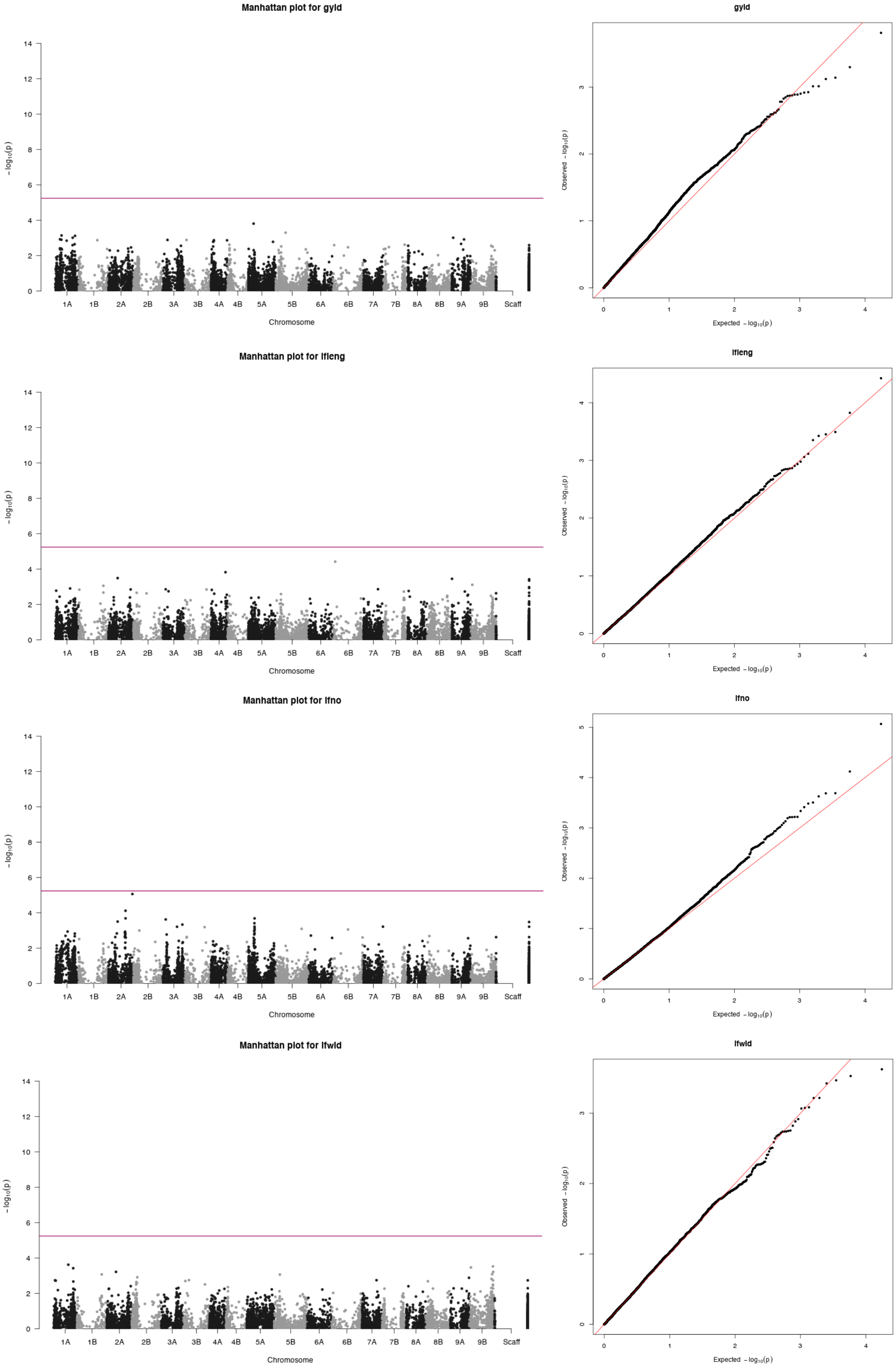

**Supplemental Figure 12.** Continued

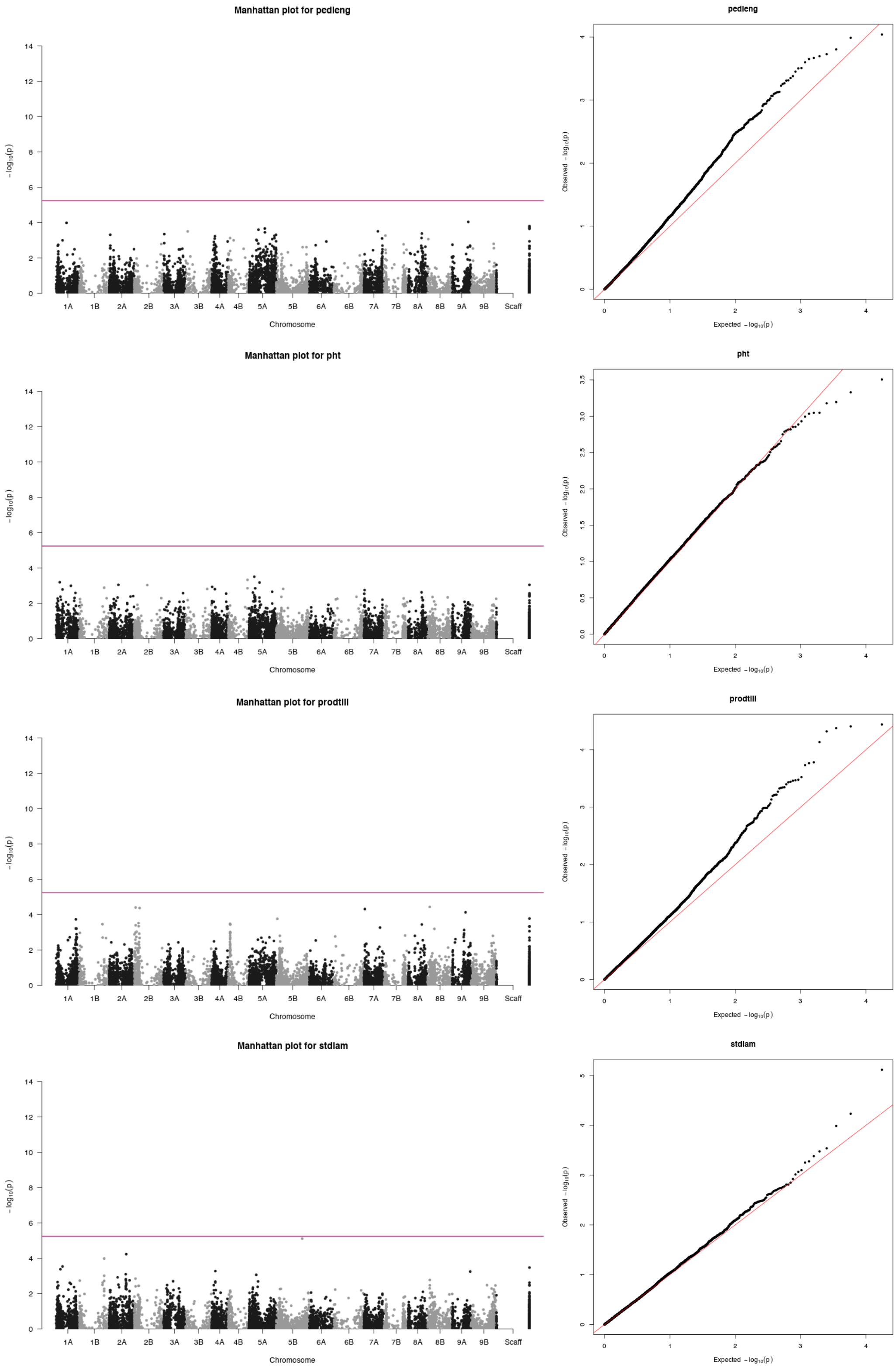

**Supplemental Figure 12.** Continued

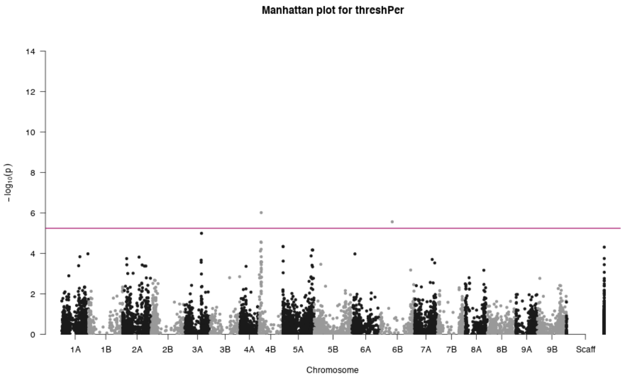


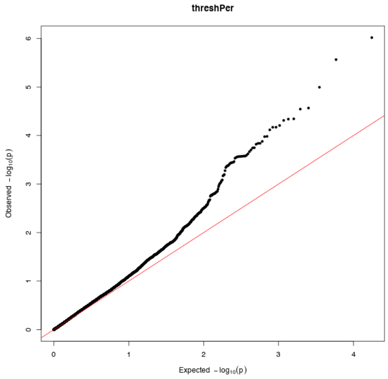

**Supplemental Figure 13. Distributions of SNP effects for 11 agronomic traits in finger millet.** The remaining two traits to total 13 in all are presented in **Fig. 5**. *stdiam,* stem diameter; *lfleng,* leaf length; *lfwid*, leaf width; *lfno,* leaf number; *fngleng,* finger length; *fngwid,* finger width; *fngno,* finger number; *pedleng,* peduncle length; *pht,* plant height; *prodtill,* production tillers; *thershPer,* threshing percentage.

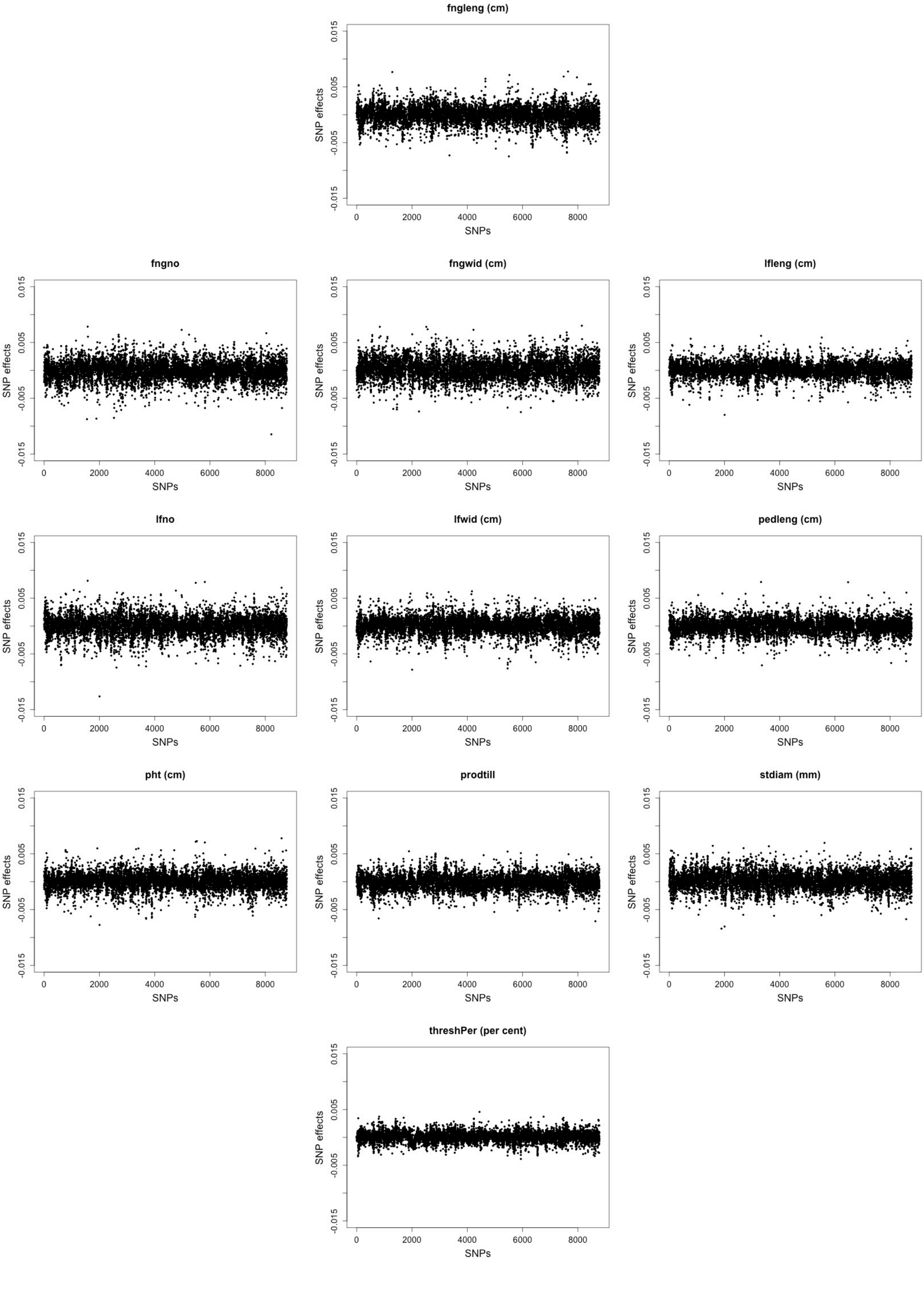
